# One-pot cloning and protein expression platform for genetic engineering

**DOI:** 10.1101/2025.08.28.672974

**Authors:** Wakana Sato, Judee Sharon, Brock Cash, Christopher Deich, Nathaniel J. Gaut, Joseph Heili, Aaron E. Engelhart, Katarzyna P. Adamala

## Abstract

In this work, we present a streamlined one-pot cloning and protein expression platform that integrates mutagenesis, plasmid assembly, and functional protein testing in a single reaction. By combining Golden Gate cloning with cell-free transcription–translation, we demonstrate efficient generation and screening of genetic variants without the need for intermediate purification or bacterial amplification. Using fluorescent proteins, luciferase enzymes, antibiotic-converting enzymes, and the violacein biosynthetic pathway, we validate the versatility of this approach for single-and multi-site mutagenesis, combinatorial variant libraries, metabolic pathway programming, and whole-plasmid assembly. By demonstrating compatibility with multiplexed reactions and multi-cistronic constructs, we establish this approach as a generalizable and automatable method for high-throughput cloning and protein engineering in synthetic biology.

## Introduction

The ability to perform rapid prototyping of large number of variants, combinations and designs of genetic circuits is driving the development of many fields of bioengineering. Those developments are driven by two main types of technical breakthroughs: the abilities to 1) clone genes and 2) rapidly test protein expression. In this work, we combine those steps into a one-pot cloning and expression pipeline.

High throughput plasmid assembly and cloning techniques can be used to assemble gene libraries and individual variants.^1^ The Golden Gate cloning, relying on Type IIS restriction enzymes^2^, have been notably one of the most commonly used techniques for those applications^3^.

On the expression testing side, while traditional live cell-based expression methods are still widely used, cell-free translation systems are rapidly gaining popularity for large scale applications.

A wide array of applications of cell-free protein expression have been demonstrated, including in biomanufacturing^4–8^, in diagnostics and biosensing^9–11^, including many successful efforts to streamline and automated development and applications^12^, and efforts towards scaling the production^13–15^.

Cell-free systems can be used in low-resource environments thanks to the ability to lyophilize the reaction mix^16^, including for educational^17,18^ and diagnostics purposes^19^, with biosensors^20^ and long term storage^21^ solutions. Cell-free systems can be automated, and are well adaptable to high-throughput regime, including investigation of product modifications^22^, enzyme and protein engineering^23–25^, and application in compartments like droplets^26,27^. Other applications include crucial role of in vitro translation in engineering synthetic cells^28^, and utility in foundational research^29^.

Cell-free systems have been used for rapid prototyping of genetic systems before^30–32^, proving the utility of this methods of protein expression for high throughput applications.

In previous applications, the cell-free expression was performed on previously synthesized or cloned DNA templates. Here we demonstrate a way to combine cloning with expression testing, combining the advantages of high throughput cloning platforms with protein function readout, eliminating the need to purify or amplify the DNA after cloning and before expression.

Performing cloning and expression reactions in the same reaction mixture demonstrates a one-pot approach to streamlined design and test of genetic elements, with broad applications for bioengineering and synthetic biology.

## Results and discussion

We developed a streamlined approach to introducing targeted mutations into a plasmid, and assembling whole plasmids, coupled with expression of the proteins encoded by that plasmid. Mutagenesis primers, encoding the new sequence, are designed to flank the target sequence, incorporating new codons and BsaI recognition sites, with additional nucleotides added to ensure efficient enzyme activity. After PCR amplification, the gene of interest is split at the mutation site, purified, and then subjected to a one-pot digestion and ligation using BsaI and T7 ligase, creating a mutated plasmid ready for expression. The process allows cloning, expression, and functional confirmation of protein variants to occur in a single reaction tube, with mutations verified through fluorescence, luciferase, or antibiotic resistance assays. (**Figure 1**).

**Figure 1.**
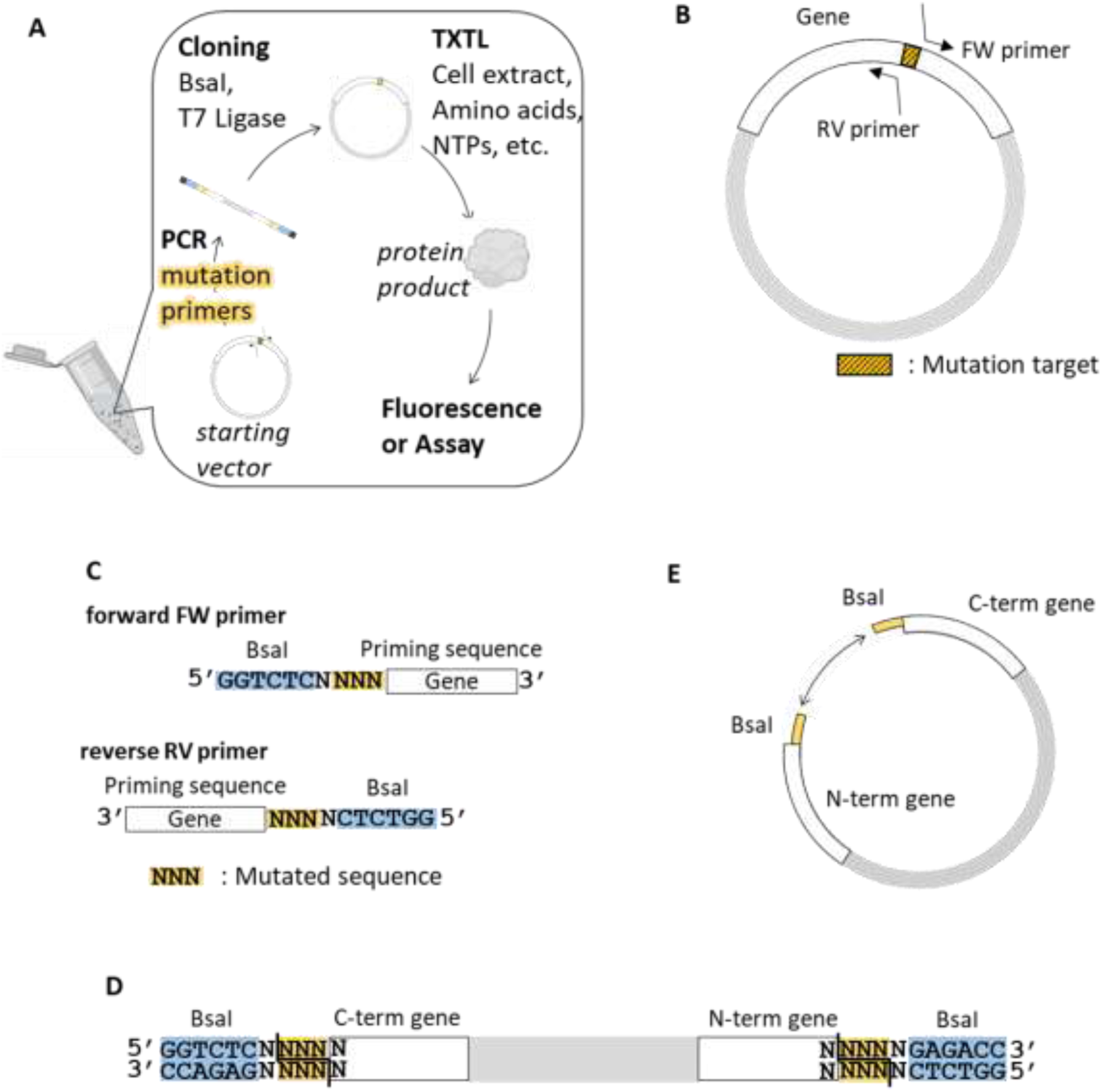
The one-pot cloning design. **A** The template plasmid before mutation. Mutation-inserting primers are designed to prime both sides of the mutation target sequence. The hatched region represents a target codon. The white region represents a gene of interest encoded in the plasmid. The light gray region represents the remaining part of the plasmid. **B** The design of a forward (FW) and reverse (RV) primers. The light blue shaded sequence is the BsaI recognition sequence. The yellow shaded three nucleotides are where replacing sequences are inserted. The white region represents parts of the gene with complementary sequences before and after (upstream and downstream) the mutation target. At the 5’ end of the primer, where is indicated as a dark gray region, we added several random nucleotides so that BsaI works appropriately on the amplified template with these primers. **C** The ends of the template after PCR. The gene of interest is split at the mutation target site. The yellow shaded three nucleotides, a new codon after cloning, are divided by a black line where BsaI cuts. **D** Schematic view of the linearized template. **E** PCR amplified linear template will be digested by BsaI, and then ligated by T7 ligase. This digestion and ligation can be processed in a one-pot reaction. This one-pot reaction is the last step of this cloning. We can express the mutated gene directly in TxTl, without any purification of the digestion and ligation reaction mixture.

Throughout this project we use transcription-translation system (called TxTl) whole cell *E. coli* lysate for protein expression.

### One-pot mutagenesis protocol validation

First, to validate the one-pot assembly method, we tested assembly of multiple colors of fluorescent proteins. Green fluorescent protein (GFP) differs from blue, cyan and yellow fluorescent proteins by three amino acids in the fluorophore (**Figure 2B**). We replaced the chromophore region of a GFP plasmid with a STOP codon, creating a plasmid expressing truncated, non-fluorescent proteins (**Figure 2A**). Then, using the one pot assembly method, we replaced the STOP codon with sequence encoding a fluorophore for one of the 4 colors of the fluorescent protein: blue BFP: Ser65, His66 and Gly67; cyan CFP: Thr65, Trp66 and Gly67; green GFP: Thr65, Tyr66 and Gly67; yellow YFP: Gly65, Tyr66 and Gly67. After mutagenesis, all samples were incubated in TxTl for 8h, and fluorescence was recorded. First, we tested three different concentrations of primers (see Materials and Methods), with 1x being the concentration typically used in the reaction. The negative control without primers recorded autofluorescence of the TxTl at the measured wavelengths. The positive control for each experiment was expression of 10nM plasmid encoding each of the fluorescent proteins. In all cases, the mutagenesis resulted in expression of the protein of the desired color, and increasing the amount of primers resulted in increased fluorescence (**Figure 2D**). Then, we tested the mutagenesis reactions with 1x primers (same primer concentration as 1x in experiments on panel **2D**), and varying the concentration of the template plasmid. Starting with 1x plasmid, the 10nM concentration used in all experiments on panel 2D, followed by twice and three times more plasmid (keeping primer concentration constant). Again, all results were compared to negative control, this time sample containing primers but no plasmid, and to positive control of the correct plasmid encoding each fluorescent protein. Unlike in the experiments increasing the primer concentration, increasing plasmid concentration did not increase fluorescence (**Figure 2E**). This indicates that the primers, not the plasmid, are the yield limiting reagent in those reactions, indicating perhaps that the least efficient step of the process is the yield of the initial PCR with the mutagenesis primers.

**Figure 2.**
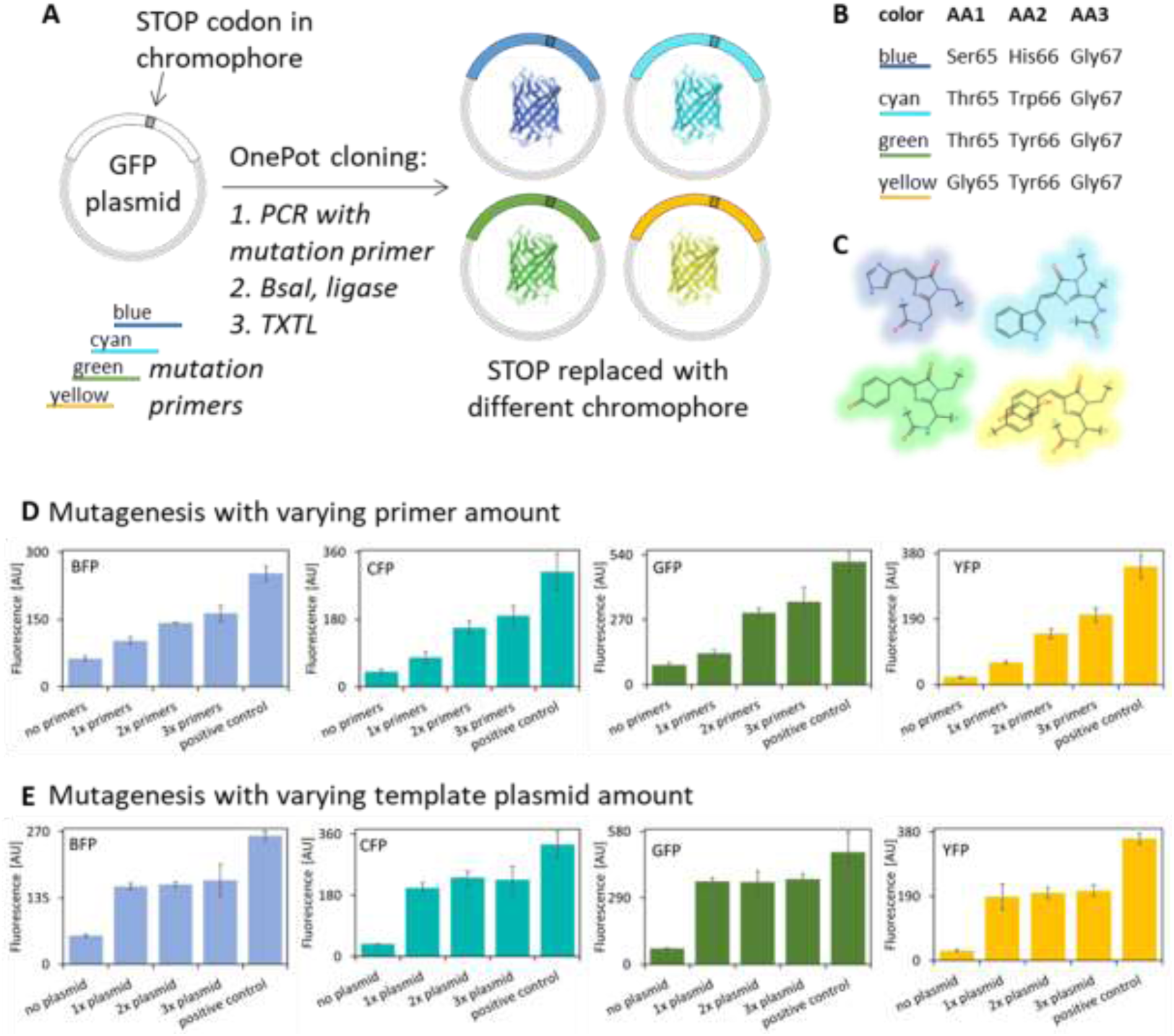
One-pot cloning mutagenesis. **A**: Fluorescent protein plasmid based on GFP sequence was prepared with a STOP codon sequence in place of the fluorescent protein chromophore. The amino acids in positions 65, 66 and 67 were removed and replaced with a STOP codon. This plasmid produces a truncated, non-fluorescent protein product. Using the one-pot mutagenesis protocol, we have re-introduced the three amino acids of the chromophore and removed STOP codon. Four different mutagenesis primers have been used, replacing the STOP codon with three amino acids encoding a different color fluorescent protein. **B**: The specific amino acids in positions 65, 66 and 67 that produce one of the 4 possible colors of a fluorescent protein. Each of the 4 primers used in mutagenesis introduced a distinct combination of those three amino acids, turning the non-fluorescent truncated protein backbone into a full length fluorescent protein of one of the 4 indicated colors. **C**: the schematic illustration of the chromophores of 4 different fluorescent proteins, created by the three amino acids indicated on panel **B**. Blue BFP: Ser65, His66 and Gly67; cyan CFP: Thr65, Trp66 and Gly67; green GFP: Thr65, Tyr66 and Gly67; yellow YFP: Gly65, Tyr66 and Gly67. **D** and **E**: One-pot mutagenesis to turn plasmid with truncated protein into a fluorescent protein. Each bar graph represents experiment with one-pot mutagenesis of the STOP-codon backbone (showed on panel A) with one of the 4 plasmids, replacing the STOP codon with three amino acids encoding a chromophore for functional fluorescent protein. Results on panel **D** show experiments where each mutagenesis was performed with constant concentration of plasmid, and increasing concentration of the mutagenesis primer. Results on panel **E** show experiments where each mutagenesis was performed with constant concentration of mutagenesis primer, and increasing concentration of the plasmid. Positive control for all graphs on panels **D** and **E**: 10nM plasmid encoding the indicated fluorescent protein, cloned via traditional methods and added directly to the cell-free reaction.

### One-pot mutagenesis to modulate enzymatic activity

Encouraged by the results with fluorescent proteins, we proceeded to mutagenize a multiple turnover enzyme and test the possibility of mutagenizing more than one site in a template in the same one-pot reaction. We chose a Firefly luciferase (PBD 1LCI), which has been previously shown to have mutant variants with increased luminescence.^33^ Some of the mutants contain two or three separate mutations (**Figure 3A**). To mutagenize one site (the variants I^423^L, D^436^G and L^530^R) we used one pair of mutagenesis primers. To introduce two mutations (in variants I^423^L+D^436^G, I^423^L+L^530^R and D436G+L530R) we used two pairs of primers, and three pairs of primers were used to create the three-point mutant (variant I^423^L+D^436^G+L^530^R). All mutagenesis experiments, including those with multiple primers, were performed in one-pot reaction, mixing all primers together. After performing the mutagenesis, followed by expression in TxTl, we analyzed the activity of the luciferase (**Figure 3B**), finding that as expected, the mutants have higher luminescence than the wild type variant. We also measured the mRNA abundance in all samples, to confirm that the difference in luminescence is primarily due to the difference in the enzymatic activity of the luciferase variants, and not due to the differences in yield of the one-pot assembly reactions (**Figure 3C**).

**Figure 3.**
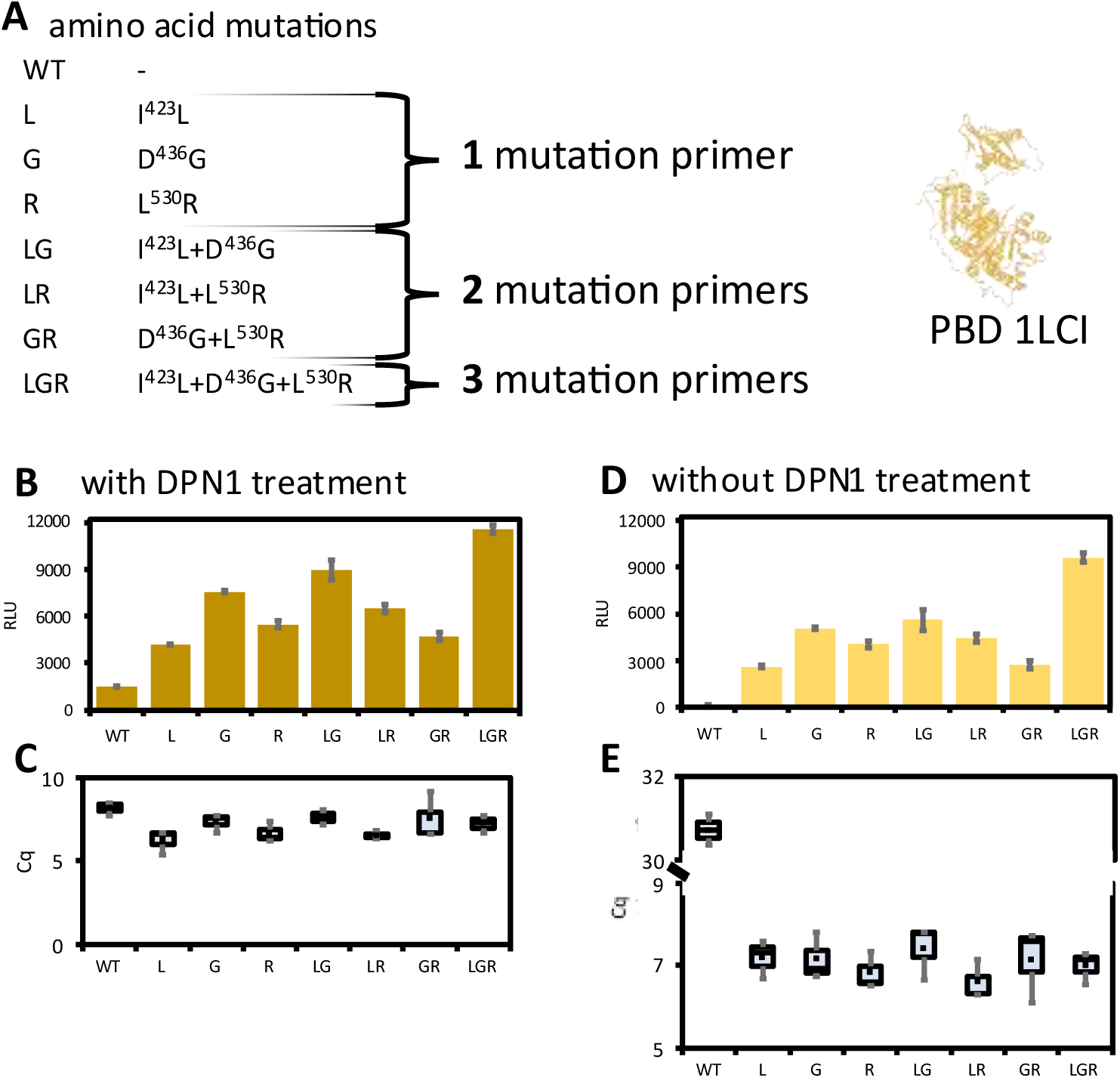
Multi-point mutations to increase enzymatic activity. **A**: Mutations in the Firefly luciferase (PBD 1LCI) lead to increased enzyme activity. **B** and **C**: One-pot mutagenesis was performed with wild type luciferase plasmid as template, and with mutagenesis primers to introduce one mutation (one set of primers, mutants L, G and R), to introduce two mutations (two sets of primers, mutants LG, LR and GR) or to introduce three mutations (three sets of primers for mutant LGR). **B**: Luminescence was measured after 8h expression in TxTl. **C**: RT-qPCR was used to measure mRNA abundance of each mutant variant. **D** and **E**: One-pot mutagenesis was performed same as in experiments on panels **B** and **C**, but omitting the DpnI enzyme digest on the plasmid after PCR. Luminescence (panel **D**) and mRNA abundance (panel **E**) was measured after 8h of TxTl reaction.

In those luciferase experiments, we suspect that the original wild type (WT) plasmid contributes to the average luminescence results, since the template originally used in the PCR remains in the reaction. To test influence of removing the original template, we performed another set of luciferase mutagenesis experiments, this time with the DpnI enzyme digest step after PCR. The DpnI enzyme digests methylated DNA, thus removing the original (bacterial amplified) template plasmid. In this case, the results from the samples containing only WT plasmid and no mutagenesis primers showed no luminescence, and all other samples showed the same ratios, but somewhat lower total fluorescence (**Figure 3D**). We confirmed those results by RT-qPCR tests, showing extremely low amount of WT mRNA, and relatively somewhat lower but similar to each other concentrations of all variant mRNAs (**Figure 3E**).

This indicated that the original WT plasmid was still present and contributing to the measured total activity. This was not an issue in the earlier fluorescent protein experiments shown on **Figure 2**, because the original template plasmid was not producing a detectable product (the STOP codon resulted in expression of truncated, non-fluorescent protein). We suggest that in mutagenesis schemes where the template plasmid has any activity on its own, a DpnI digest step is added after PCR to remove the template plasmid.

Next we tested the one-pot assembly method on an enzyme where mutations change not only enzymatic efficiency, but also substrate specificity. *Acremonium chrysogenum* deacetoxy/deacetylcephalosporin C synthase (acDAOC/DACS) catalyzes the ring-expansion of penicillin N to deacetoxy-cephalosporin C (DAOC) and hydroxylates DAOC to deacetylcephalosporin C (DAC). The enzyme has a broad substrate specificity, converts substrates besides penicillin N, such as penicillin G^34,35^. Wu et al. identified the R308 residue of acDAOC/DACS affects its substrate specificity and catalytic activity by saturated mutagenesis at position R308, where is predicted to control the interaction with its substrate^34,35^. This finding led us to believe that acDAOC/DACS can be a good testing platform for our one-pot cloning technique, to demonstrate the ability to modify enzymatic activity by single amino acid mutation. We first confirmed acDAOC/DACS, expressed in TxTl from isolated plasmids, converts penicillin analogs in our system using HPLC **(Figure 4B)**. Then we performed one-pot mutagenesis of acDAOC/DACS to create variants at position R308, and we performed activity assays over several penicillin analogs **(Figure 4C)**. We created 19 mutants, and each demonstrated different conversion rates of the 4 penicillin variants tested (penicillin G, penicillin V, ampicillin and carbenicillin).

**Figure 4.**
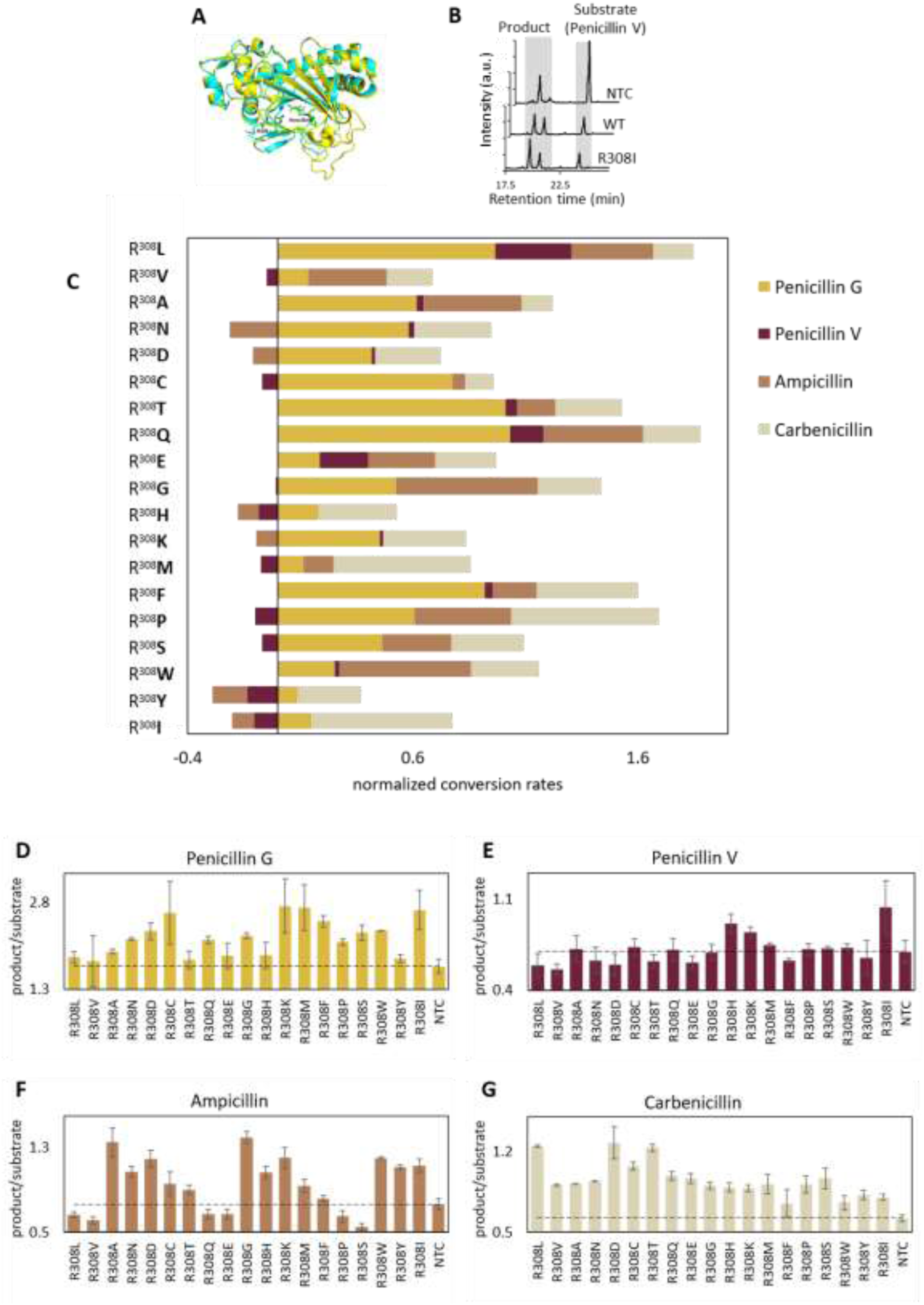
One-pot mutagenesis of antibiotics converting enzyme mutants. **A**: Predicted structures of DAOC/DACS, a bifunctional enzyme involved in biosynthesis of antibiotics from *Acremonium chrysogenum* (yellow) and DAOCS from *Streptomyces clavuligerus* (cyan) in complex with penicillin G. Penicillin G and the R308 residue were shown as sticks, atoms are colored: C-green, N-blue, O-red, S-yellow. **B**: An example of HPLC chromatogram for analysis of penicillin analogue conversion. The ratio between substrate peak and product peaks changed between reactions. NTC: No template control. WT: Wild type enzyme. R308I: Mutant where arginine on position 308 was replaced with isoleucine. More examples of the HPLC chromatograms are on Figure S1. **C**: One-pot cloning and expression strategy was used to create acDAOC/DACS mutants with the point mutation indicated on the Y axis. Penicillin analogs conversion rates are shown aggregated for all penicillin analogues for each mutant. The assay data were normalized with no template control. All mutant sequences are listed in **Table S1**. **D** – **G**: Individual data for penicillin analogue conversion with all mutants and NTC (no template control). The dotted line is a visual reference representing NTC levels. Error bars represent SEM, n=3. Western Blot analysis of expression of acDAOC/DACS in TxTl is on Figure S2. Calibration curves for HPLC antibiotic detection are on Figure S3.

### Generation of combinatorial pools

Encouraged by this versality of the one-pot cloning scheme, we decided to expand on the fluorescent protein cloning scheme shown on **Figure 2**, and take advantage of the ability to perform multiple point mutation experiments in one sample shown on **Figure 3**, to create combinatorial sets of reactions with different protein product ratios. One again we started with the fluorescent protein backbone with chromophore replacing the STOP codon. We created a well plate design where each well contained the same amount of template, and the same total amount of mutagenesis primers, but the primers were encoding different combinations of protein variants. Each well contained some combination of primers encoding conversion of the template to blue, cyan, green or yellow fluorescent protein (**Figure 5A**). We designed the experiment to test different ratios of primers resulting in different ratios of fluorescent protein products. The design of the experiment is shown on **Figure 5B**, **5C**, **5D** and **5E**, with panels showing the relative amount of each primer pair encoding the indicated fluorescent proteins. Each well contained either none, 1x, 2x, 3x or 4x of each of the color primer pair, to the total of 4x primers in each well. The mutagenesis and expression was performed in well plate format.

**Figure 5.**
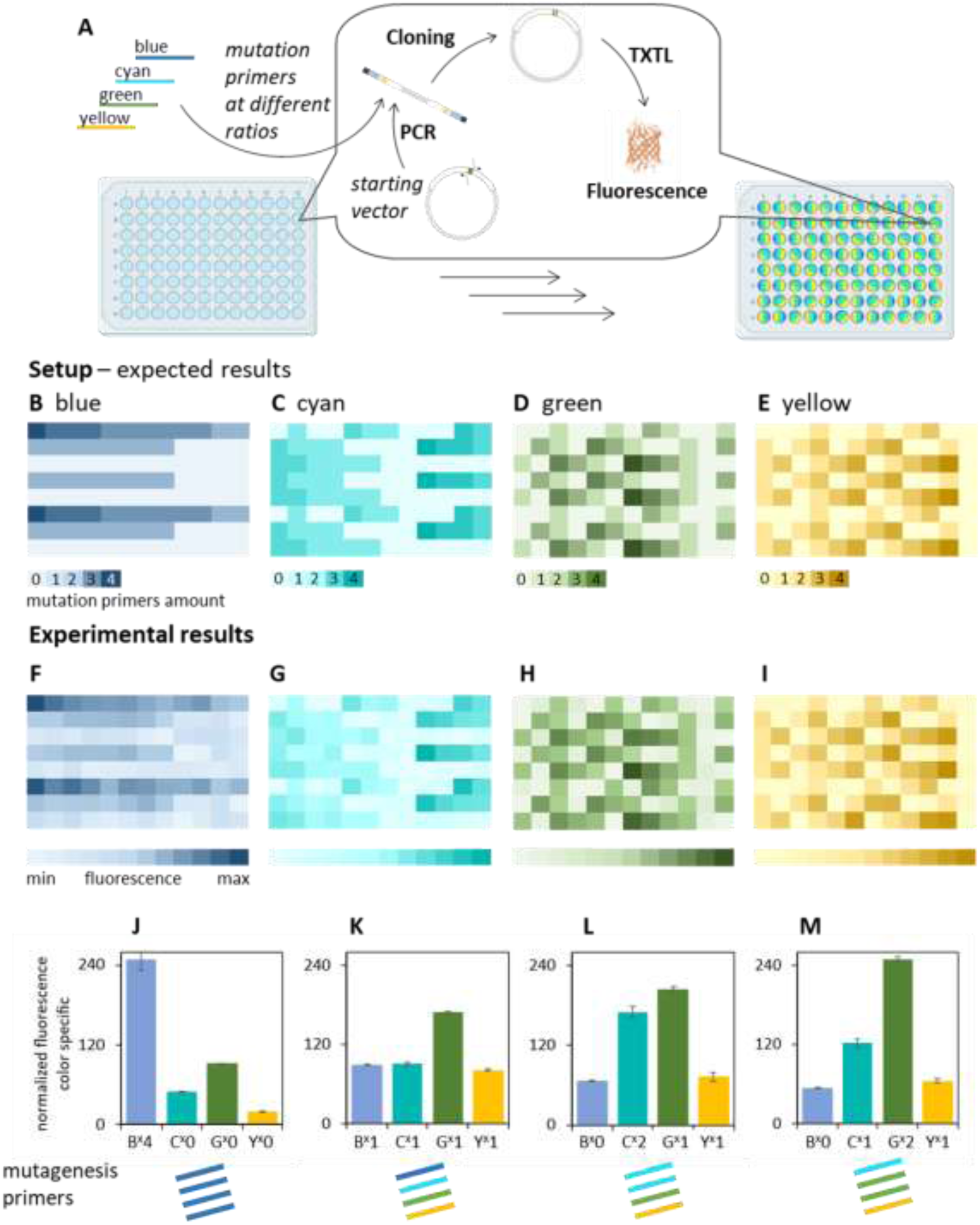
One-pot plasmid assembly in high throughput format. A: Starting vector, eGFP plasmid with STOP codon mutation in the first amino acid position in the chromophore, was subjected to one pot cloning protocol with one of four mutation primers. Each of the mutation primers encoded sequence of the chromophore for one of the four possible colors of fluorescent proteins based on the GFP architecture. Those primers, and resulting fluorescent proteins, were: Blue BFP - Ser65, His66 and Gly67; cyan CFP - Thr65, Trp66 and Gly67; green GFP - Thr65, Tyr66 and Gly67; yellow YFP - Gly65, Tyr66 and Gly67. After one pot cloning, resulting plasmids were expressed in cell-free translation reaction. **B-M:** All reactions were set in 96-well plate format. Each well contained a different ratio of mutagenesis primers, with the total DNA concentration constant. Panels b-e show the theoretical setup of each experiment: each well of the plate contained a different ratio of each of the 4 primers, between 0 (no primer) to 4 (all primer in this well). The total concentration of DNA template and primers was constant in all wells. Each primer is shown on separate panel (**B**: primers for blue BFP, **C**: cyan CFP, **D**: green GFP, **E**: yellow YFP). Panels **F-I** show fluorescent measurements results from experimental setup shown on panels **b-e**, after cloning procedure and 12h incubation in cell-free translation reaction. Each panel shows fluorescence measured in separate channel, with excitation and emission matching the target fluorescent protein (f: blue, g: cyan, h: green and i: yellow). Panels **J-M** show individual data for 4 selected wells, representing different experimental conditions. Panel **J**: well A1, containing 4x blue primer (denoted B^x^4) and no other primers (denoted C^x^0, G^x^0 and Y^x^0). Panel **K**: well G3, containing 1x of each primer (denoted B^x^1, C^x^1, G^x^1 and Y^x^1). Panel **L**: well H1, containing no blue primer (denoted B^x^0), 2x of cyan primer (denoted C^x^2), and 1x green and yellow primer (denoted G^x^1 and Y^x^1). Panel **M**: well H4, containing no blue primer (denoted B^x^0), 2x of green primer (denoted G^x^2) and 1x of cyan and yellow primers (denoted C^x^1 and Y^x^1). Individual data points for all values shown on panels **F**, **G**, **H** and **I** are on figures S4, S5, S6 and S7. Schematic of the setup of the well plate experiments is on Figure S8.

After mutagenesis and TxTl incubation, we measured fluorescence from the plate in all 4 fluorescent color channels. The results (Figure **5F**, **5G**, **5H** and **5I**) correspond well to the expected results. In wells with higher amount of the primer for a particular color, more fluorescence of that color protein was observed.

This confirms that one-pot assembly and expression pipeline can be used to design large throughput, combinatorial pools of gene variants.

### Programming of enzymatic pathway

The violacetic acid pathway is a bacterial pathway producing pigment molecules^36^, with all enzymes of that pathway previously expressed in cell-free reactions^37^. The products and intermediates of that pathway are easily identifiable on HPLC, and the pathway output can be modulated depending on the identity of the enzymes used in each particular reaction. (**Figure 6A**) The reagents in the pathway are: TRP, L-tryptophan; IPAI, indole-3-pyruvic acid imine; IPAID, IPAI dimer; CPA, chromopyrrolic acid; PDV, prodeoxyviolacein; DV, deoxyviolacein; PV, proviolacein; VIO, violacein.

**Figure 6.**
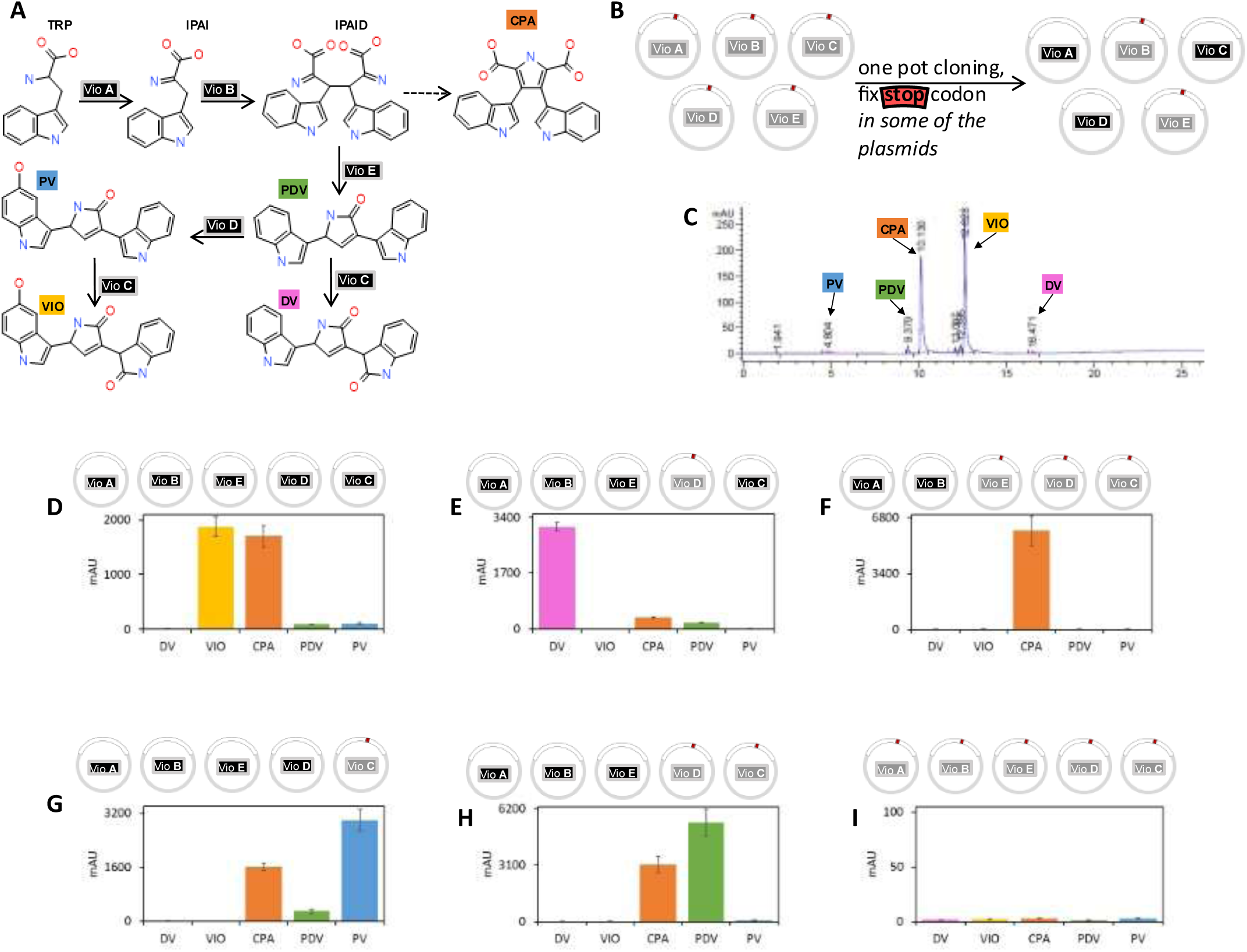
One-pot reactions for the assembly of metabolic engineering pathway. **A**: The schematic of the violacein biosynthesis pathway, with small molecule products, intermediates and substrates, and enzymes catalyzing specific steps. The output of the pathway can be controlled by the identity of the enzymes in the reaction mix. B: The schematic of one-pot cloning reaction to program the Vio pathway. All Vio enzymes have been modified to introduce a STOP codon at position 50, resulting in truncated and inactive products. One-pot cloning reactions are used to fix (by removing STOP codons) some of the enzymes, creating a pool with a combination of active enzymes programming the pathway. **C**: An example of HPLC analysis of Vio reaction products, separating all the products and intermediates. **D** – **I**: Different Vio biosynthesis pathways built by one-pot mutagenesis of different plasmids, with all the mutagenesis done on a pool of plasmids mixed together. Each mutagenesis removed the STOP codon from the selected plasmids, enabling expression of a full length functional enzyme. After cloning and expression of the enzyme in TxTl for 12h, the reactions were analyzed on HPLC. **D**: mutagenesis of all plasmids, creating a complete Vio pathway with all enzymes. **E**: mutagenesis of all plasmids except VioD. **F**: mutagenesis of only VioA and VioB. **G**: mutagenesis of all plasmids except VioC. **H**: mutagenesis of all plasmids except VioD and VioC. **I**: negative control, no mutagenesis plasmids were used, so all Vio genes remained inactivated by the premature STOP codon.

We modified all plasmids of the Vio pathway to insert a STOP codon at position 50, resulting in production of non-functional truncated enzymes from each plasmid. To rescue each enzyme, we designed primer sets for the one-pot mutagenesis (**Figure 6B**). We performed the mutagenesis on samples containing all 5 Vio pathway enzyme plasmids in each reaction, and a varying set of mutagenesis primers designed to rescue a particular combination of enzymes. All reactions were incubated in TxTl and analyzed on HPLC. The product distribution from each experiment corresponds to the expected product distribution for each of the combinations of the enzymes (**Figure 6D**, **6E**, **6F**, **6G** and **6H**). The negative control, incubation of the template plasmids without any mutagenesis primers, results in no detectable products (**Figure 6I**).

Those experiments demonstrate how one-pot assembly technique can be used to combinatorially program a multi-enzyme metabolic engineering pathway.

### Combinatorial assembly of whole plasmids

Introducing mutations and rescuing sub-sets of plasmids is useful in bioengineering and design-test-learn cycle of synthetic biology. In addition to that, sometimes there is a need for larger scale genome rearrangements, including combinatorial construction of whole new plasmids. We tested whether the one-pot assembly protocol can be useful in such cases, to assemble a whole plasmid, not just mutagenize part of a gene.

To perform plasmid assembly experiments, we prepared linear templates containing different variants of the promoter sequence, open reading frame (ORF) gene insert, antibiotic resistance cassette, and a bacterial origin of replication (ORI). We then designed primers for PCR introducing the BsaI overhangs necessary for the one-pot assembly. We performed one pot assembly reactions, in each case producing a complete vector, which was then tested in TxTl reaction (**Figure 7A**).

**Figure 7.**
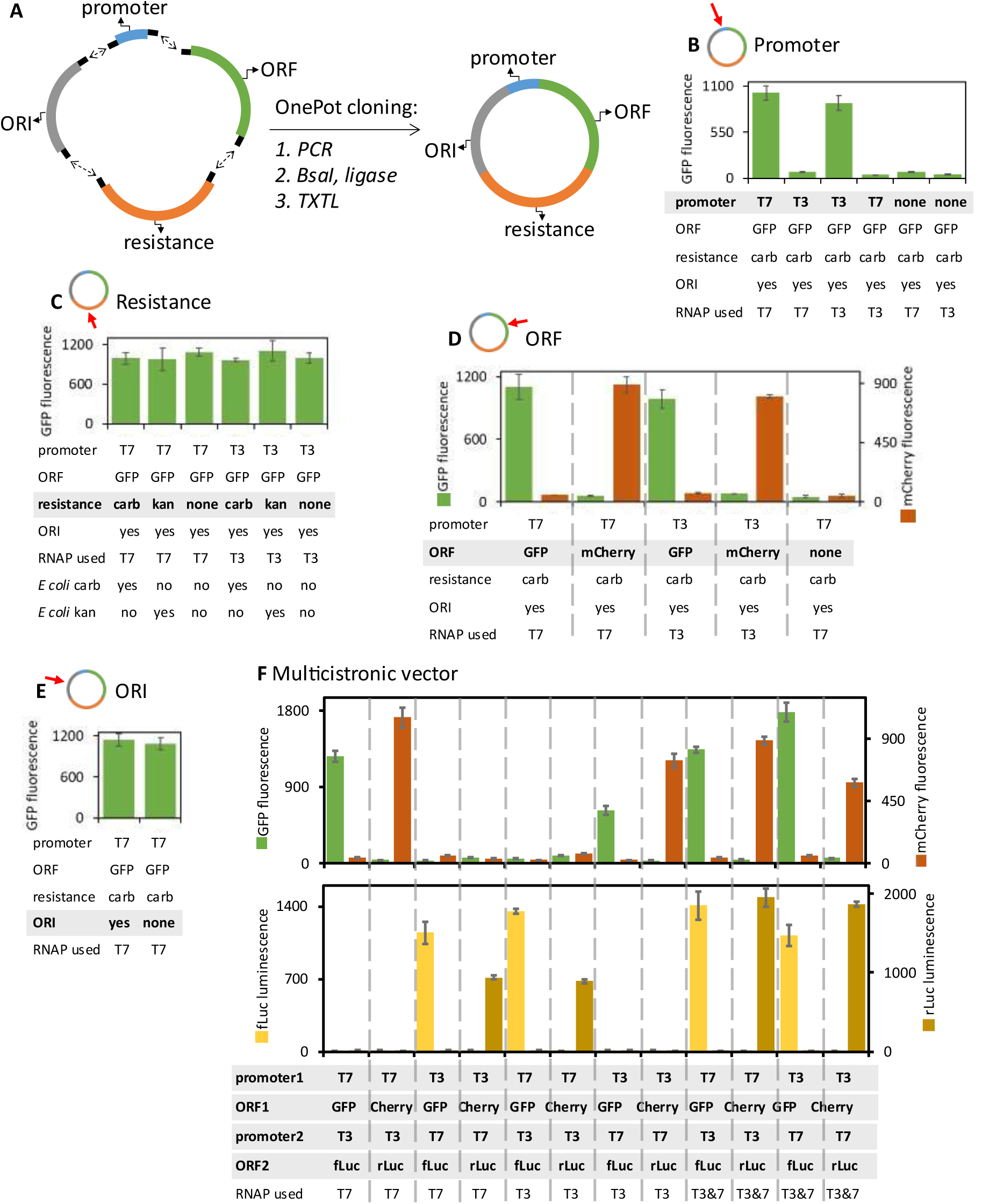
Complete plasmid assembly in one-pot reaction. **A**: The one-pot cloning was used to assemble a whole vector, including promoter, open reading frame (ORF), antibiotic resistance, and origin of replication (ORI) regions. **B** – **E**: Assembly of vectors with one of the regions varied in each experiment. In all experiments, mutagenesis was followed by 8h TxTl incubation and analysis. **B**: Assembly of vectors with a different promoter. Vectors were built with T7 or T3 RNA polymerase promoter, or without a promoter region, and expression tests were performed in TxTl reaction mixture containing either T7 or &3 RNA polymerase. **C**: Assembly of vectors with beta-lactamase gene (carbenicillin resistance) or KanR gene (kanamycin resistance). TxTl expression tests were performed, and part of the mixture was also used to transform live *E coli* cells, to test the antibiotic resistance. The transformed cells were grown either on carb or kan plates overnight, and monitored for colony forming (yes means colonies were detected, no means no colonies after 24h). **D**: Assembly of vectors with different open reading frame. The gene inserted into the ORF was either sequence encoding GFP or mCherry fluorescent proteins. During the analysis after expression, fluorescence was recorded for both GFP and mCherry channels. **E**: Assembly of a vector with or without the origin of replication (ORI). **F**: The one-pot assembly of multi-cistronic vectors. Two promoter and two ORF regions were assembled for each vector. The first ORF was either GFP or mCherry, the second ORF was either firefly luciferase or Renilla luciferase. After cloning and 8h TxTl expression, fluorescence was recorded from all samples, and then samples were analyzed in luciferases activity assays, measuring activity of both firefly and Renilla luciferases.

We assembled plasmids with different promoter, either T7 or T3 RNA polymerase promoter, or without a promoter region (in that case, the BsaI assembly sites were flanking a non-coding, non-binding random region of DNA). Those promoter variant plasmids were tested in TxTl with different RNA polymerase, either T7 or T3 RNA polymerase in the reaction. Fluorescence was only detected from samples where the intended assembled promoter matched the RNA polymerase used in the reaction, so a T7 promoter assembly in reaction with T7 RNAP, and T3 promoter assembly in reaction with T3 polymerase (**Figure 7B**).

Next we assembled plasmids with different antibiotic resistance. We used either beta-lactamase gene (carbenicillin resistance) or KanR gene (kanamycin resistance) cassettes, or no antibiotic resistance (the BsaI assembly sites were flanking a non-coding, non-binding random region of DNA). Those vectors were assembled with either T7 or T3 resistance. Assembled plasmids were tested in TxTl, and as expected, all plasmids produced fluorescent signal, since antibiotic resistance is not needed for plasmid performance in TxTl (**Figure 7C**). To test whether the assembled plasmids indeed have the indicated resistance, each sample was used for transformation of live *E coli*, which were then plated on plates containing either carbenicillin or kanamycin. After 24h at 37C, colonies were found only on the plates where the cloned antibiotic resistance matched the antibiotic used on the plate: carb plasmids produced colonies on carb plate, kan plasmids produced colonies on kan plates, and plasmids without resistance produced no colonies on either plate.

Next, we assembled vectors with a different open reading frame (ORF), using inserts encoding GFP or mCherry fluorescent proteins. After cloning and incubation in TxTl, we tested fluorescence in both green and red channel for each sample. Fluorescence detection results followed the expected pattern: samples with GFP gene insert produced green fluorescence, and samples with mCherry insert produced red fluorescence (**Figure 7D**).

The next set of experiments assembled vectors with and without bacterial origin of replication. The ORI has no effect on TxTl expression, so as expected, both vectors with and without ORI produced fluorescent signal in TxTl (**Figure 7E**). To confirm the assembly of the ORI region, samples were transformed into live *E coli*, plated on carb plates and incubated for 24h at 37C. Colonies were only found on the plate with bacteria transformed with plasmid that did contain ORI.

Finally, we investigated the possibility of assembling a multi-cistronic vector, with more than one open reading frame. We prepared fragments for two promoter regions (T7 and T3), and two ORF variants: the first ORF was either GFP or mCherry gene, the second ORF was either firefly luciferase or Renilla luciferase. After one-pot assembly and TxTl incubation, we analyzed fluorescence in both green and red channel and analyzed luciferase activity in both Firefly and Renilla assays. Both the fluorescence and luminescence analysis followed the pattern expected from the experimental design. Green fluorescence was found in samples with vector assembled with GFP insert and promoter matching the RNA polymerase used in TxTl, red fluorescence also was only found in samples with mCherry insert and promoter matching the polymerase used.

Similarly, firefly and Renilla luciferase activities were only detected in samples where the enzyme gene and promoter matched the luciferase reagent and RNA polymerase used **Figure 7F**). This confirmed that a multi-cistronic plasmid can be assembled and expressed using the one-pot protocol.

### Conclusions and future outlook

Here we demonstrated a way to couple cloning reaction with protein expression, in a streamlined one pot format compatible with well-plates and future automation. We tested the assembly and expression of several types of enzymes and reporter proteins, and demonstrated multi-point mutagenesis and assembly of multi-part vectors.

The two main limitations of this system are: the need to eliminate BsaI sites from any other part of the assembled vector (a constraint shared by all Golden Gate based cloning systems^38^), and the compatibility of the protein product with cell-free expression. While *in vitro* translation in bacterial lysate is capable of installing many post-translational modifications^22^, some still remain difficult^39^.

The work presented in this paper focuses on assembly of plasmids, but since bacterial amplification is not needed to test expression, this method could also be applied to any DNA fragments with promoter, gene sequence and terminator. Some cell-free systems can express linear templates, eliminating the need for circular plasmids^40–42^.

Some possible future adaptations of this system could include using PURE^43,44^ instead of TxTl, to combine the versatile cloning strategy with the genetic code expansion tools^45^. Other possibilities include using this system in automation platform using microfluidics^46,47^ and directed evolution^48^, to take advantage of the high throughput assembly and testing potential.

## Acknowledgments

We thank Dr. Vincent Noireaux for the gift of T7 RNAP plasmid and for helpful discussions about troubleshooting and optimizing the cell-free protein expression system. We thank Dr. Claudia Schmidt-Dannert for the gift of Vio pathway plasmid and for helpful discussions about optimizing the pathway. This work was supported by the BioMADE award S-PC04-P-72, the Alfred P. Sloan Foundation grant G-2024-22710, and the NSF Award 2419641 (to K.P.A), and NIH R01 GM152459 (to A.E.E).

## Materials and Methods

The below cloning protocols describe an example procedure for one of the plasmids used in this work, to provide detailed description of an example of the one-pot protocol and other procedures used in this work.

### Traditional cloning

All the PCR reaction for this cloning was performed using the recommended PCR protocol provided at NEB website (https://www.neb.com/). acDAOC/DACS WT and R308I genes were obtained as gBlocks™ Gene Fragments from IDT. The two genes (acDAOC/DACS-WT and acDAOC/DACS-R308I) were PCR amplified with primers (Fprimer-restriction-cloning and Rprimer-restriction-cloning) and Q5® High-Fidelity 2X Master Mix (M0492S, NEB). After PCR, the amplified genes were agarose gel purified and digested with with AgeI-HF (R3552L, NEB) and MluI-HF (R3198S, NEB) in 50 μl reaction containing 1x CutSmart® Buffer (NEB, B7204S). After digestion, genes were purified using GenCatch^TM^ PCR Cleanup Kit (Epoch, 2360250).

The backbone plasmid (empty-vector-plasmid) was also digested with AgeI-HF and MluI-HF, then agarose gel purified. The digested backbone was dephosphorylated using Shrimp Alkaline Phosphatase (rSAP) (NEB, M0371L). The acDAOC/DACS genes and the backbone were ligated using T4 DNA Ligase (NEB, M0202L), and successfully cloned plasmids were selected on LB agarose plates containing 100 μg/ml Carbenicillin (Goldbio, C-103-50). The sequences were confirmed with seq-primer-1 and seq-primer-2 using sequence service offered by MCLAB (https://www.mclab.com/).

### Linearization PCR

PCR amplification was performed to make individual linearized plasmids. The 50 μl of PCR reactions containing 1ng of the plasmid (acDAOC/DACS-WT-plasmid), 0.4 μM of forward primer, 0.4 μM of reverse primer, and 1x of LongAmp® Taq 2X Master Mix (NEB, M0287S) was incubated on a thermal cycler (BIO-RAD T100 Thermal Cycler, 186-1096) at the NEB recommended PCR condition: Initial denaturation at 94°C for 30sec; denaturation at 94°C for 20 sec, annealing at 52°C for 15 sec, and extension at 65°C for 195 sec, which this cycle was repeated 30 times; final extension at 65°C for 10 min; hold at 4°C until further treatment.

This mixture was used for the next step of the one-pot reaction, or in the initial optimization steps the PCR product could be analyzed by electrophoresis.

To analyze the PCR reactions (not part of the one-pot protocol) After the PCR reaction, 50 μl of PCR product was mixed with 10 μl of DNA loading dye (NEB, B7024S) and loaded on agarose gel electrophoresis performed with 1% agarose gel containing 1x sybr safe (Thermo fisher science, S33102) at 125 V for 45 min in TAE buffer (40 mM Tris (pH 7.6), 20 mM acetic acid, 1 mM EDTA). The 3940 bp PCR products were purified from the agarose gels with a gel extraction kit (Epoch life science, 2260250), and the yields were measured with nanodrop (Thermo fisher science, ND-1000 Spectrophotometer).

### Restriction enzyme digestion and ligation setup

Restriction enzyme digestion and ligation were performed in the 40 μl of reaction mixture containing 25 ng/μl of PCR product, 1.55 units/μl of BsaI-HF®v2 (NEB, R3733L), 150 units/μl of T7 ligase (NEB, M0318L), 1mM ATP, 1x CutSmart® Buffer (NEB, B7204S). The incubation was performed on the thermal cycler: Initial warming up at 37°C for 3 sec; digestion at 37°C for 5 min and ligation at 20°C for 5 min, which this cycle was repeated for 25 times; heat inactivation at 80°C for 20 min; hold at 4 °C until further treatment. The completed one-pot cloning mixtures were used for transformation or TXTL.

### Transformation from the one-pot cloning mixture

To isolate mutant acDAOC/DACS plasmids, the ligated samples were transformed into homemade chemically competent BL21(DE3) cells prepared using Mix & Go! *E.coli* Transformation Buffer Set (Zymo, T3002) and successfully cloned plasmids were selected on LB agar plate containing 100 μg/ml carbenicillin. The mutation was confirmed by MCLAB sequencing service with primers of seq-primer-1 and seq-primer-2.

### TxTl

70 μl of TxTl reactions were prepared to express the mutant proteins. TxTl reaction mixture: 5 nM plasmid (acDAOC/DACS-WT-plasmid, acDAOC/DACS-R308I-plasmid) or 21.8 μl of one-pot reaction, 12 mM of magnesium glutamate, 140 mM of potassium glutamate, 1 mM of Dithiothreitol (DTT), 1x energy mix, 1x amino acid mix, 0.8 U/μl of RNase inhibitor (NEB, M0314L), 1 μM of T7 RNA polymerase (purified by ourselves, and 1x cell-free extract). The TXTL reactions were incubated at 30°C for 8 hours and held at 4°C until being used for the assay. As a control, empty-vector-plasmid was used.

10x Energy mix: 500 mM HEPES pH 8.0, 15 mM of ATP, 15 mM of GTP, 9 mM of CTP, 9mM of UTP, 2 mg/μl of *E. coli* tRNA mixture, 0.68 mM of folinic acid, 2.6 mM of coenzyme-A, 15 mM of spermidine, 40 mM of sodium oxalate, 7.5 mM of cAMP, and 300 mM of 3-Phosphoglyceric acid (3-PGA).

10x Amino acids mix (pH 6.5): 400 mM KOH and 20 mM of each amino acids (alanine, arginine, asparagine, aspartic acid, cysteine, glutamic acid, glutamine, glycine, histidine, isoleucine, leucine, lysine, methionine, phenylalanine, proline, serine, threonine, tryptophan, tyrosine, and valine).

### Western blotting

7.5% resolving gels were prepared by mixing 4.61 ml water, 1.78 ml 40% Acrylamide, 0.95 ml 2% Bis-Acrylamide, 2.5 ml 1.5M Tris-HCl (pH 8.8), 100 μl 10% SDS, 50 μl 10% Ammonium Persulfate (APS), and 5 μl Tetramethylethylenediamine (TEMED). 4% stacking gels were prepared by mixing 2.82 ml water, 0.38 ml 40% Acrylamide, 0.20 ml 2% Bis-Acrylamide, 1 ml 1.5M Tris-HCl (pH 8.8), 40 μl 10% SDS, 20 μl 10% Ammonium Persulfate (APS), and 2 μl Tetramethylethylenediamine (TEMED).

20 μl TxTl reaction mixture was mixed with 20 μl 2x sample loading dye (100 mM Tris HCl, 2.5% SDS, 20% Glycerol, 4% Beta mercaptoethanol, 0.1% Bromophenol Blue) and heat-denatured at 95°C for 5 min. The 40 μl of mixed samples were loaded in gel wells. 5 μl Blue stain protein ladder (Gold Biotechnology, P008-500) was also loaded. The gels were run at 100V for 1 hour in 1x SDS running buffer (25mM Tris base, 200mM Glycine, 1% SDS). The proteins on the gel were transferred to the nitrocellulose membrane at 100V for 1 hour in 1x transfer buffer (25mM Tris base, 200mM Glycine). After transfer, the membranes were incubated in 20 ml of 5% milk in TBST (20mM Tris, 150mM NaCl, 0.05% tween) on a rocker for 1 hour. Then 4 μl of anti-His antibody [BioLegend, 652505] was added and incubated at 4°C on a rocker overnight. The next day, the membrane was washed with TBST three times, then incubated with TBST for 10min three times. After the wash, the membrane was incubated in 4 μl of HRP Goat anti-rat IgG antibody (BioLegend, 405405) in 20ml of 5% milk for 1 hour. The membrane was washed with TBST three times, then incubated with TBST for 10min three times. The chemiluminescence reaction was performed using SuperSignal West Pico PLUS Chemiluminescent Substrate (Thermo Fisher Scientific, 34577), and gels were visualized with a gel imager (Bio-Rad, Universal Hood III).

### Antibiotic conversion assay

The antibiotic degradation activity was measured *in vitro* assay. The final concentration of the reagents in the reaction mixture are as follows: 50mM MOPS (pH 7.5), 1.8 mM FeSO4, 4 mM of ascorbate, 2.56 mM of α-ketoglutarate, 28 mM penicillin analogs (Penicillin V, Amoxicillin, or Carbenicillin). For the 30 μl of reaction preparation, MOPS (pH 7.5), 1.8 mM FeSO4, ascorbate, and water were pre-mixed and aliquoted into 3 μl of TXTL (one-tenth volume of entire reaction volume) tubes that had expressed acDAOC/DACSs. Then, α-ketoglutarate was added to the individual tube. Finally, a penicillin analog was added and incubated at 30 °C for 20 min. After the incubation, 30 μl (the equal volume of reaction volume) of ethanol was added to stop the reaction, and the sample was dried with a speedbag. The dried pellet was reconstituted with 60 μl of water and used for HPLC analysis.

### HPLC analysis

The HPLC analysis was performed with Agilent ZORBAX Bonus-RP 3.5 µm 4.6 x 150 mm (Agilent, 863668-901) and the HPLC system (On-line vacuum degasser, Hewlett-Packard, G 1322A; Autosampler, Hewlett-Packard, G 1313A; Column Compartment, Hewlett-Packard, G 1316A; Diode Array Detector, Agilent G1315B; Quaternary Pump, Agilent, G1311A). HPLC gradient condition was as follows: solvent A; H2O+ 0.1% TFA, Solvent B; ACN + 0.1% TFA, 0-5min 5%B; 5-35min 5-65%B; 35-40min 65-95%B; 40-50min 95%B; 50-51min 95-5%B; 51-60min 5%B, 40 °C. 2 μl of sample was injected and detected the UV absorbance at 220 nm. The enzyme activity was determined by taking the ratio of the peak area of the substrates and products. The peak representation was determined by comparing the HPLC chromatograms between enzyme-expressed reaction and reaction with empty vector (NTC; No Template Control). The presented values were calculated with the formula: (The sum of product peak area/ The sum of substrate peak area)-(The sum of background product peak area/The sum of background substrate peak area). Calibration curves for antibiotic detection are on Figure yyS7.

## Supplementary materials

**Figure S1.**
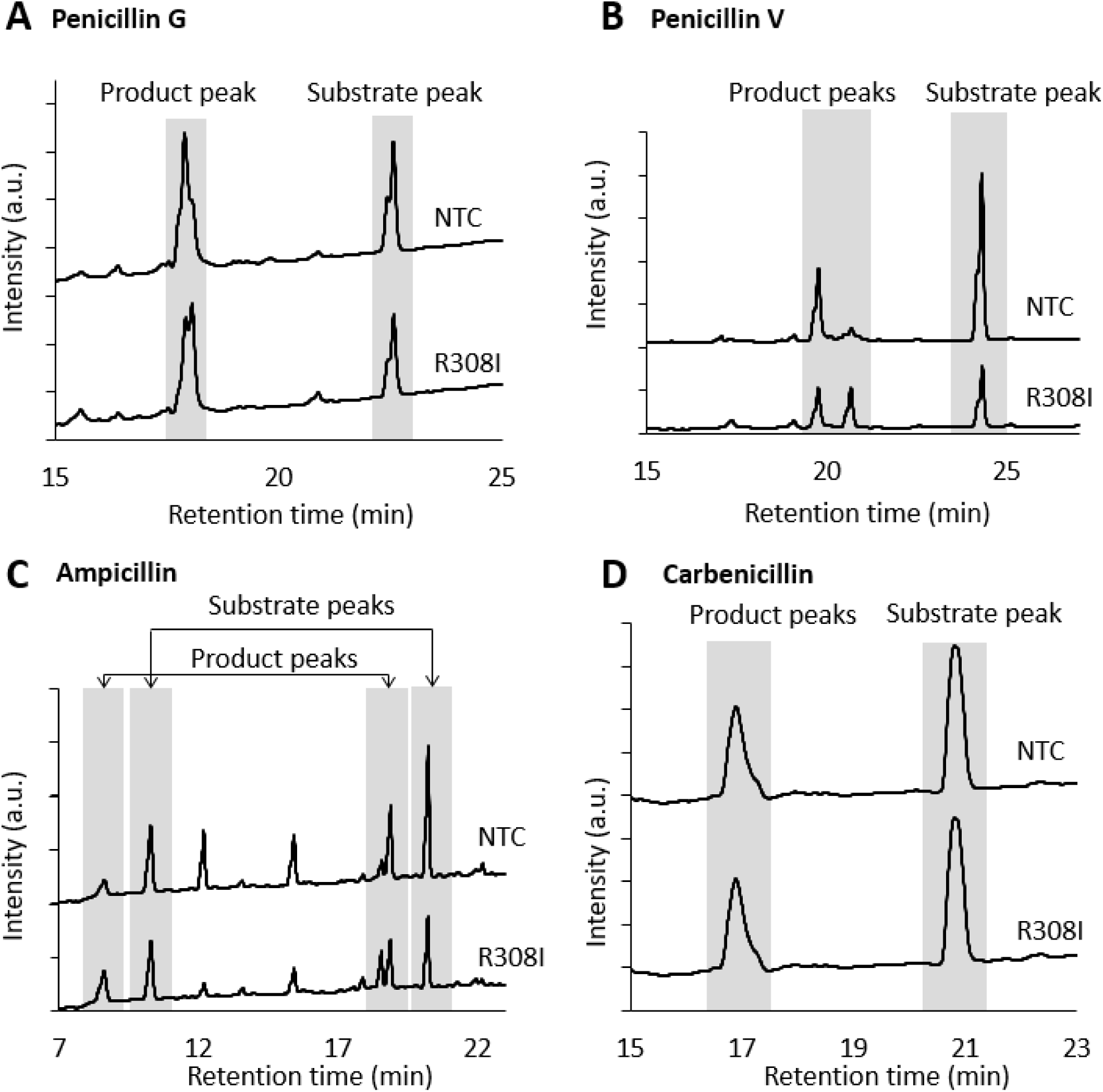
HPLC chromatogram examples. The substrate and product peaks were determined by comparing to a chromatogram performed without substrates (Data not shown). The HPLC chromatograms represent the catalytic activities of acDAOC/DACS mutant (R308I) expressed from one-pot cloning reactions. (A) Penicillin G, (B) Penicillin V, (C) Ampicillin, and (D) Carbenicillin were used as its substrates.

**Figure S2.**
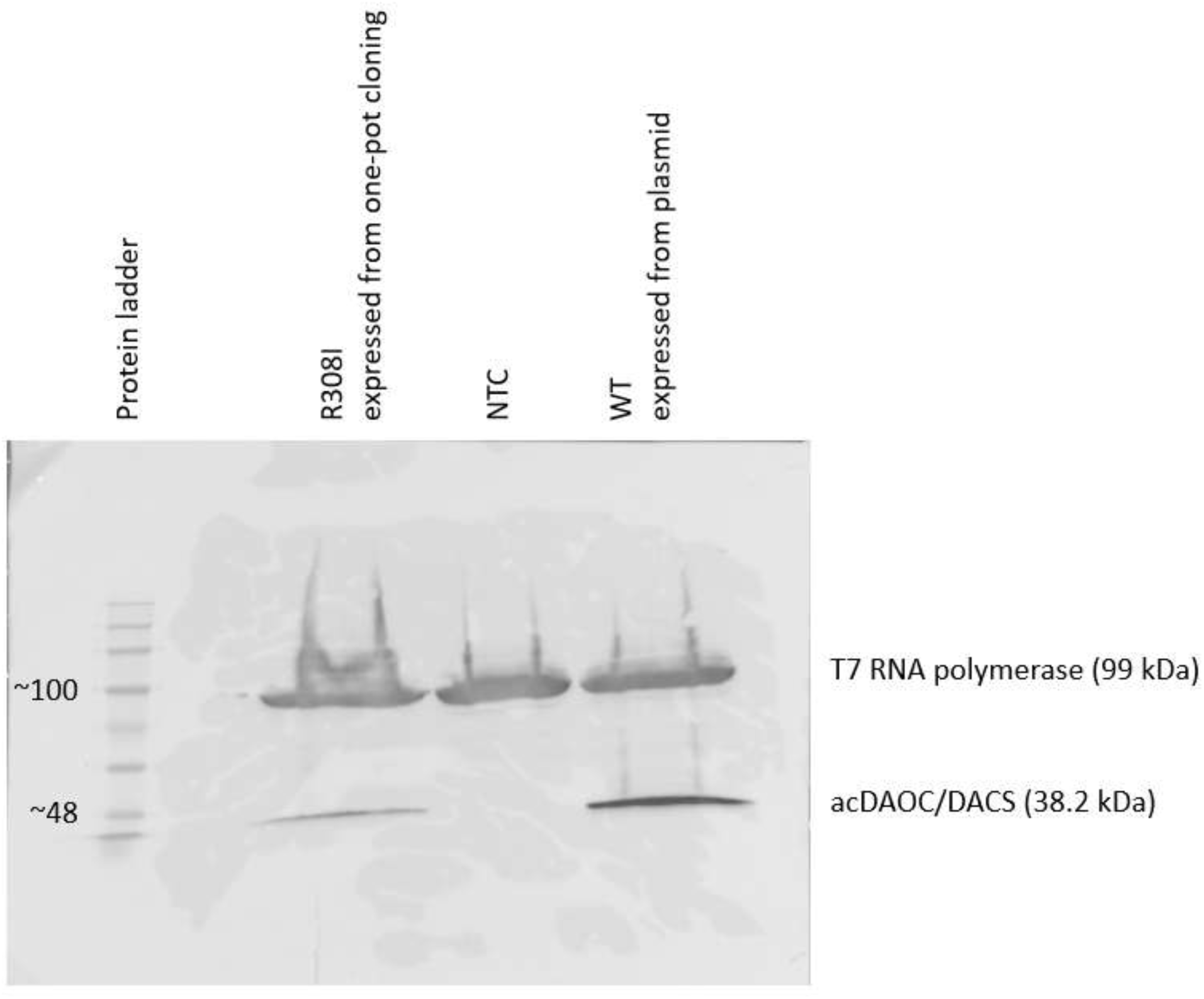
acDAOC/DACS expressions in TXTL. The representative western blotting gel. 5 μl of Blue stain protein ladder was loaded in the left well. The samples loaded were TXTL with R308I one-pot cloninig mixture, no template control (NTC), or wild type (WT) plasmid (from left to right). The thick bands around 99 kDa are C-term His-tagged T7 RNA polymerases. The bands around 38 kDa are C-term His-tagged acDAOC/DACS. The acDAOC/DACS expressed better with purified plasmid than one-pot cloning mixture.

**Figure S3.**
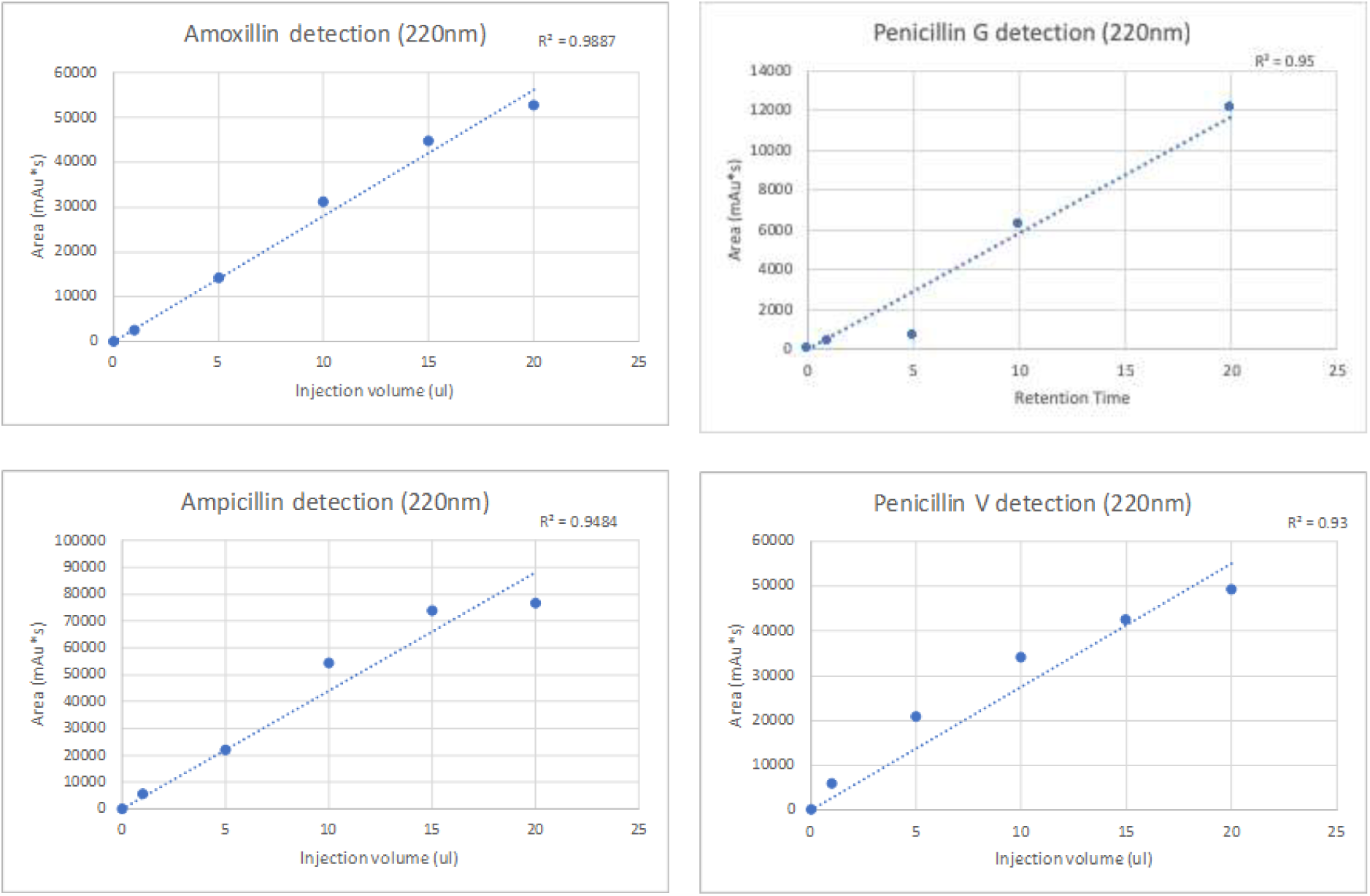
Calibration curves for HPLC antibiotic detection.

**Figure S4.**
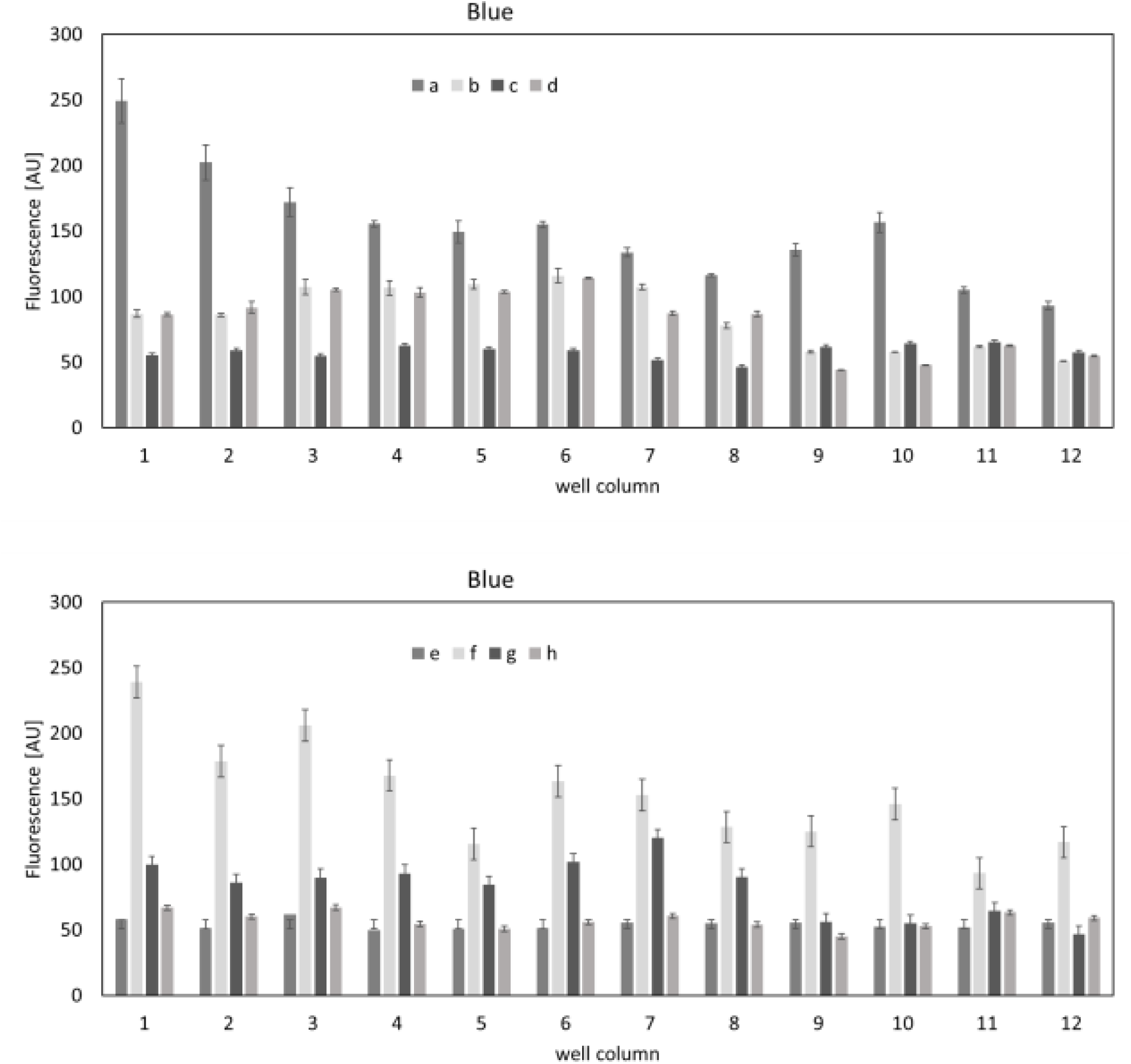
Individual data points for heat map on figure 5. The blue protein fluorescence measured in each well of the 96-well plate, with well column indicated by the X graph axis and well row indicated by the legend (rows a, b, c and d on first graph and rows e, f, g and h on second graph). Error bars are S.E.M., n=3.

**Figure S5.**
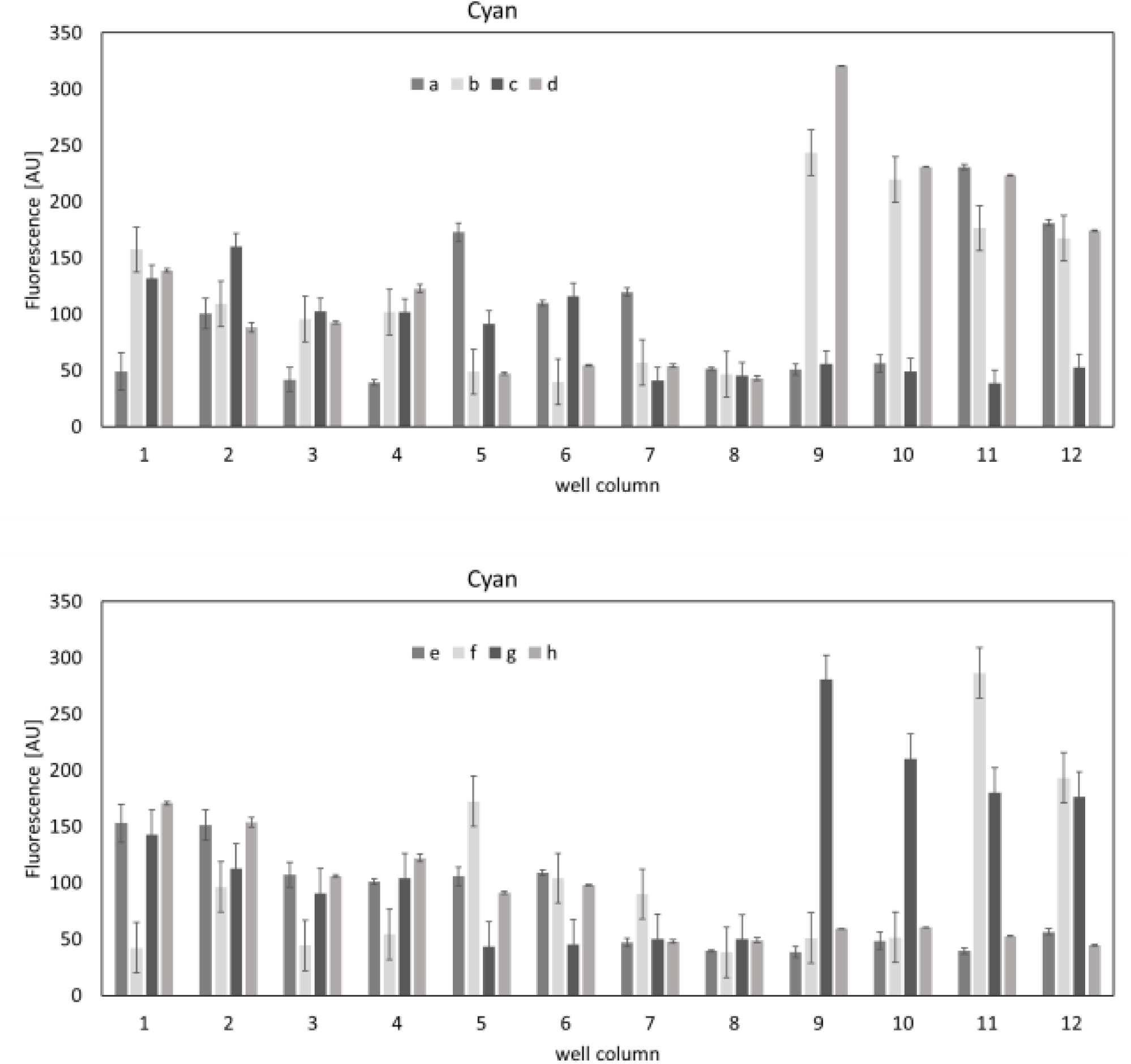
Individual data points for heat map on figure 5. The cyan protein fluorescence measured in each well of the 96-well plate, with well column indicated by the X graph axis and well row indicated by the legend (rows a, b, c and d on first graph and rows e, f, g and h on second graph). Error bars are S.E.M., n=3.

**Figure S6.**
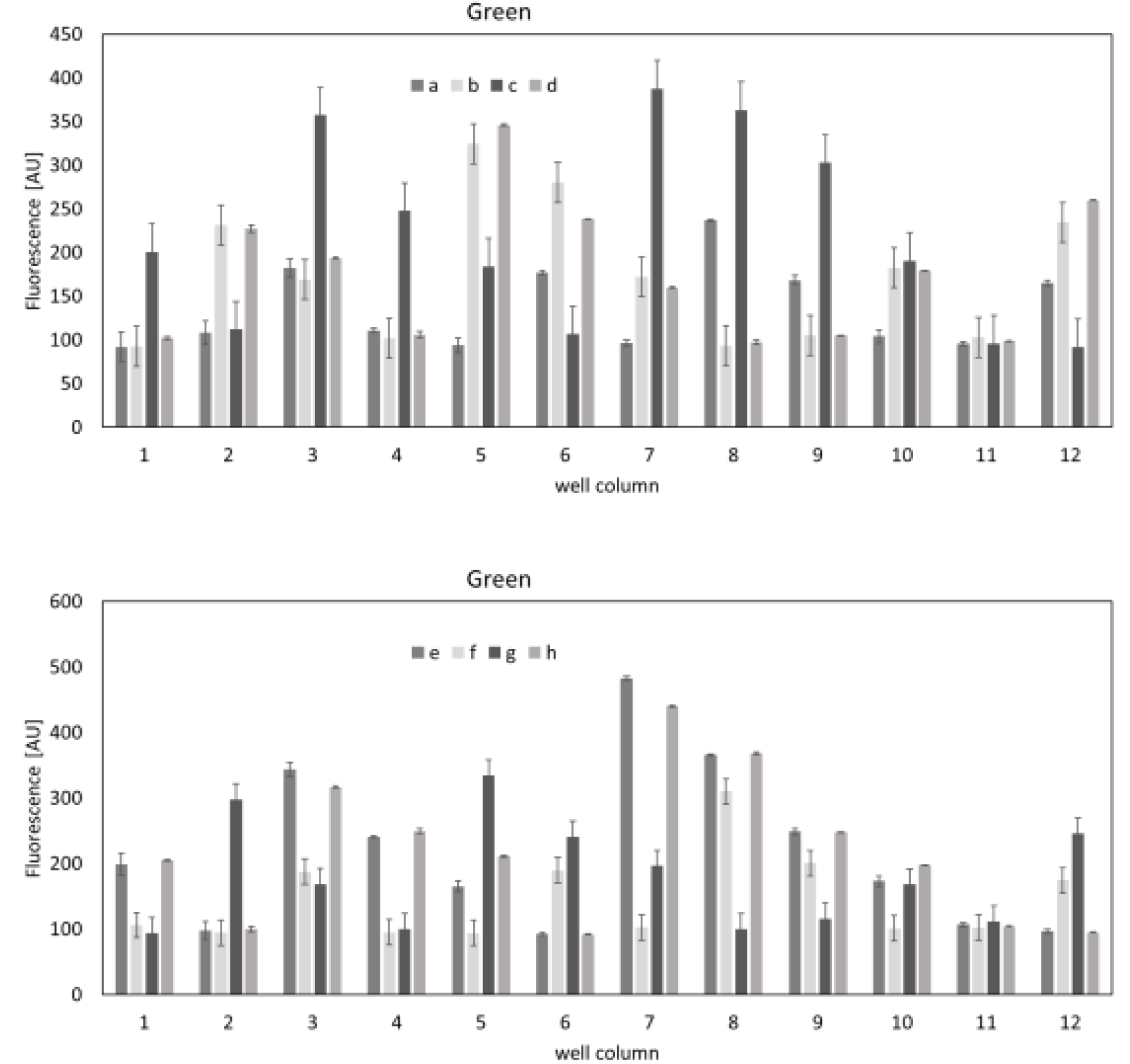
Individual data points for heat map on figure 5. The green protein fluorescence measured in each well of the 96-well plate, with well column indicated by the X graph axis and well row indicated by the legend (rows a, b, c and d on first graph and rows e, f, g and h on second graph). Error bars are S.E.M., n=3.

**Figure S7.**
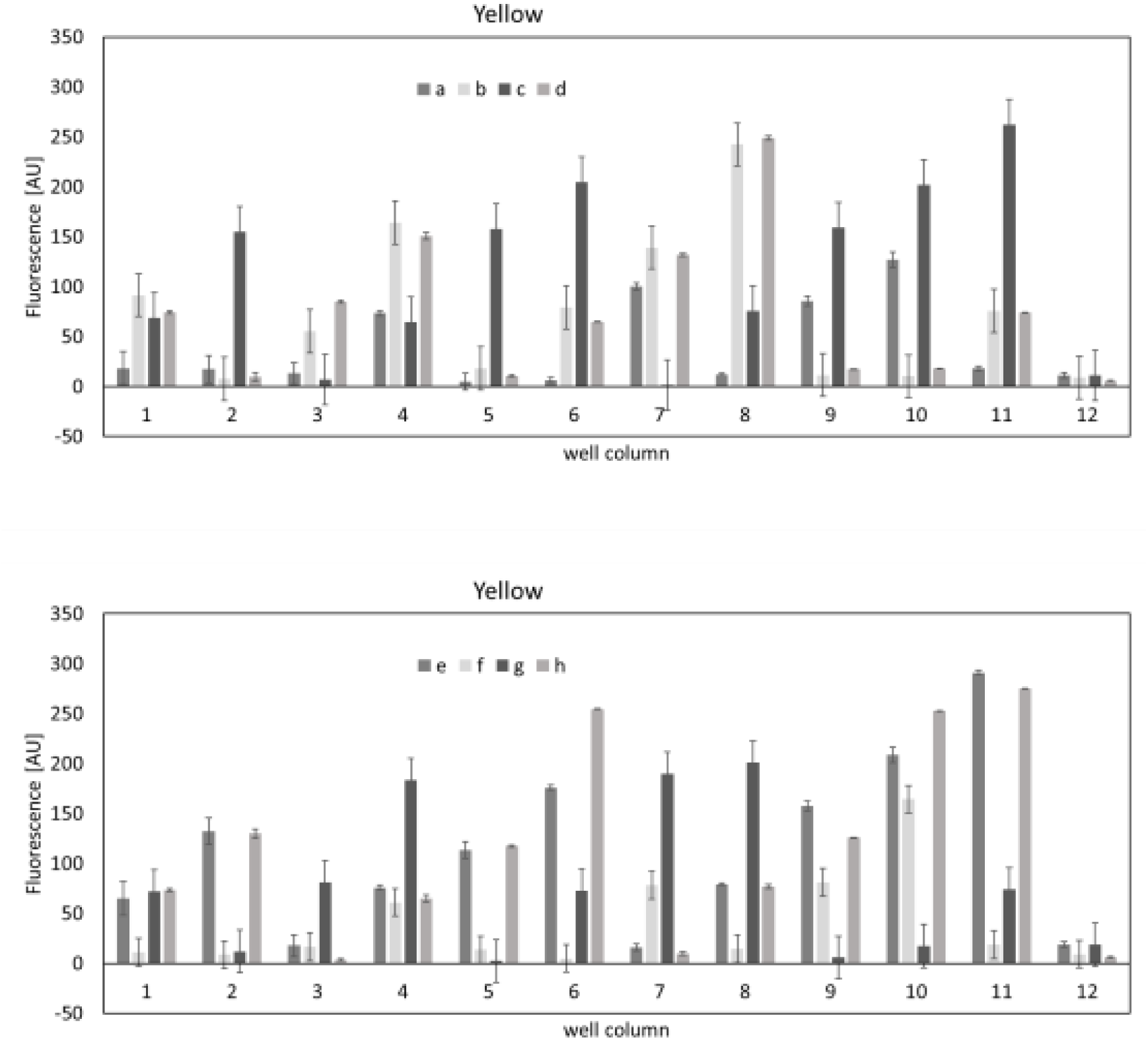
Individual data points for heat map on figure 5. The yellow protein fluorescence measured in each well of the 96-well plate, with well column indicated by the X graph axis and well row indicated by the legend (rows a, b, c and d on first graph and rows e, f, g and h on second graph). Error bars are S.E.M., n=3.

**Figure S8.**
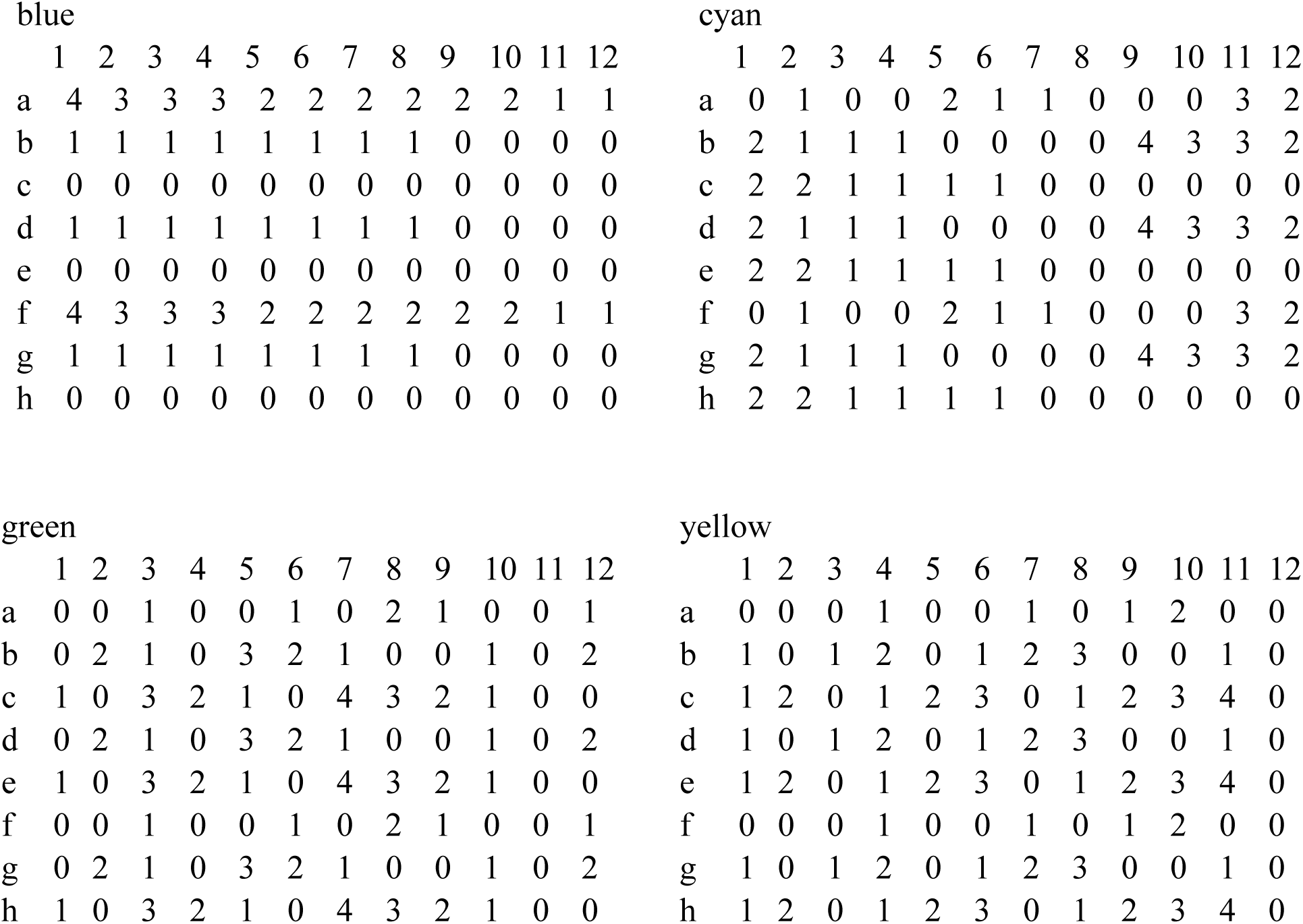
The setup of the well-plate experiments for fluorescent protein color change in one-pot reaction. The schematics show the amount of primer for each color in each well of a 96-well plate experiment. 0 – no mutagenesis primer for this color. 1, 2, 3 and 4 – increasing amount of mutagenesis primer.

---

**Table S1.**
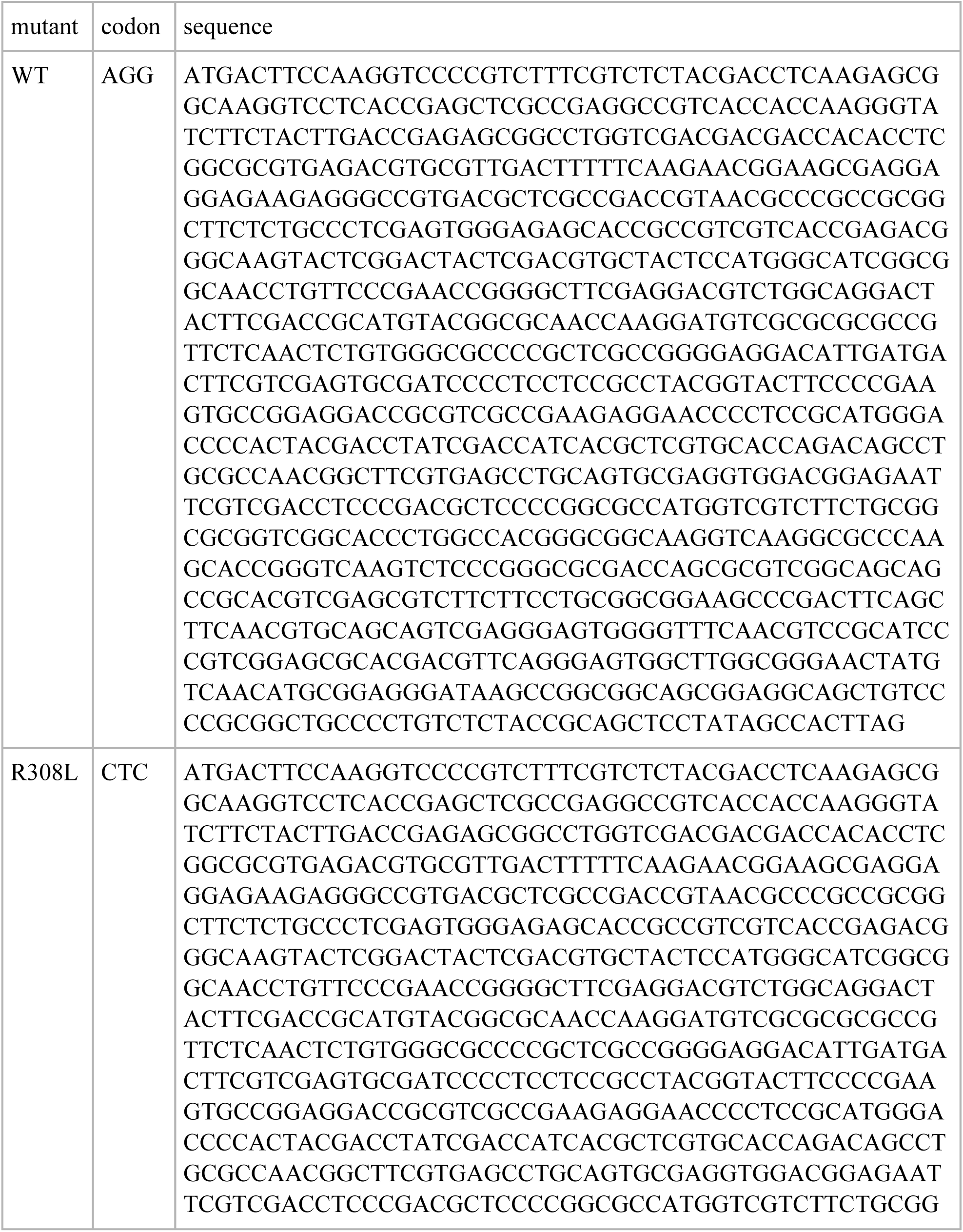
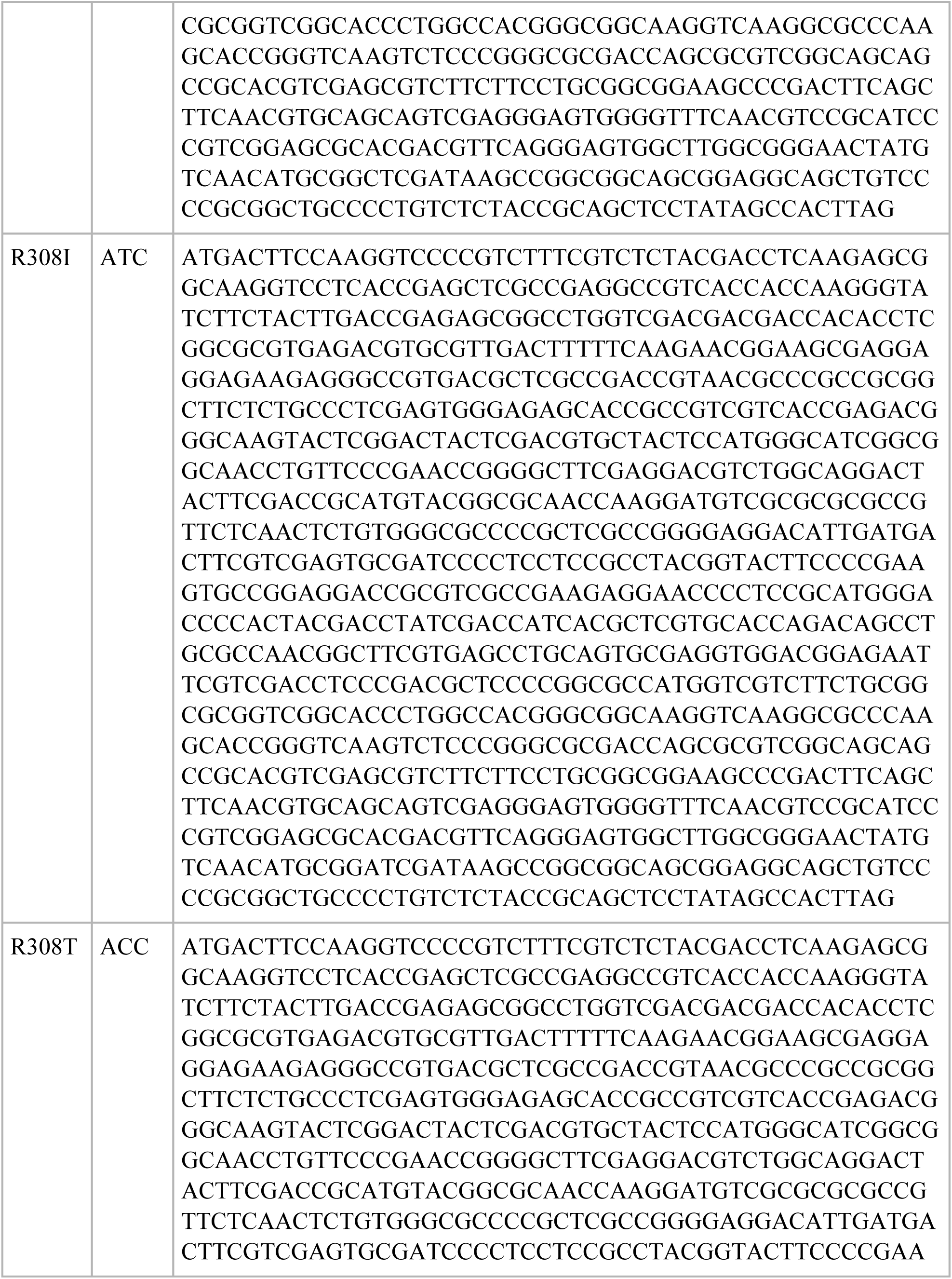
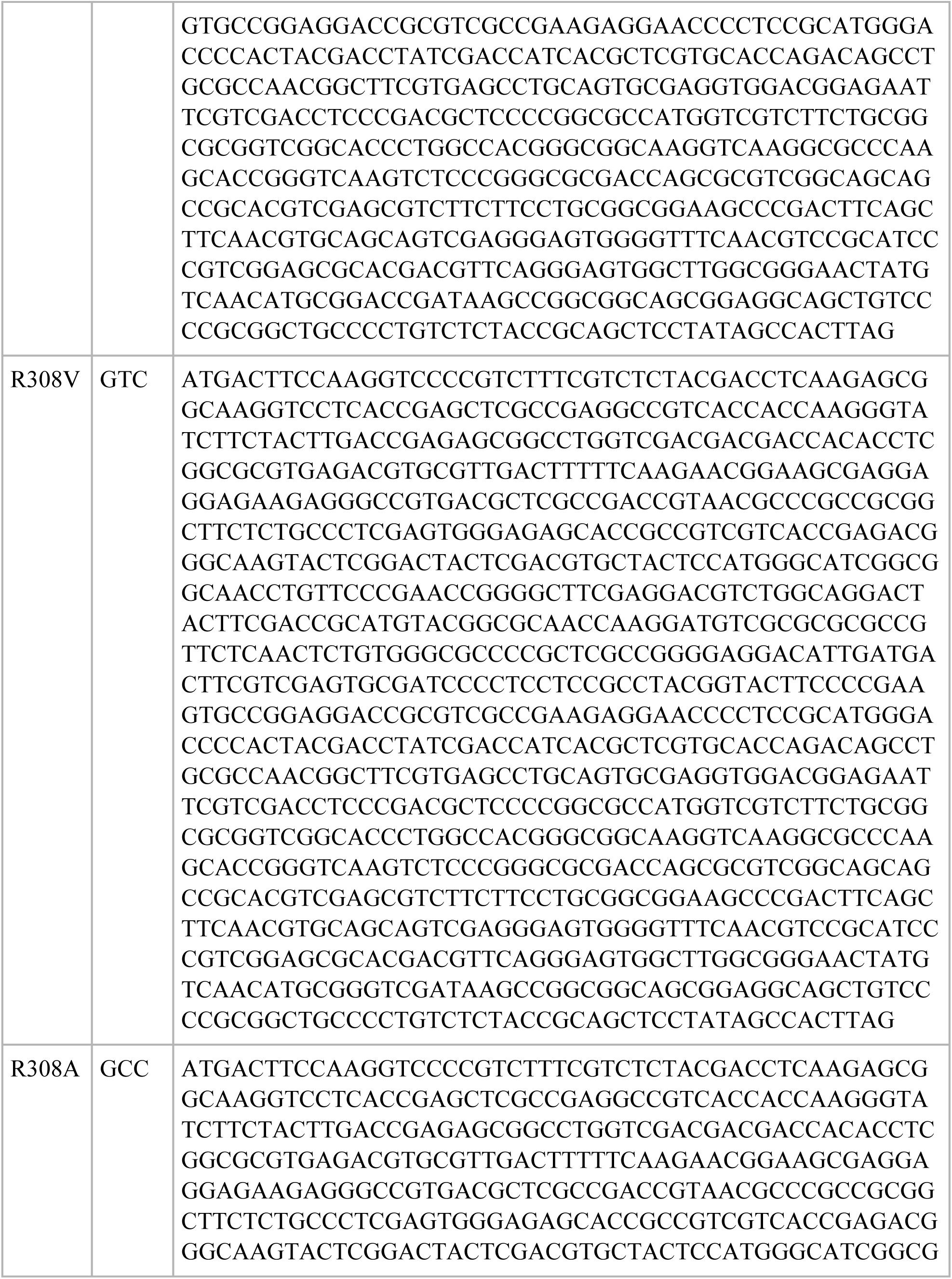
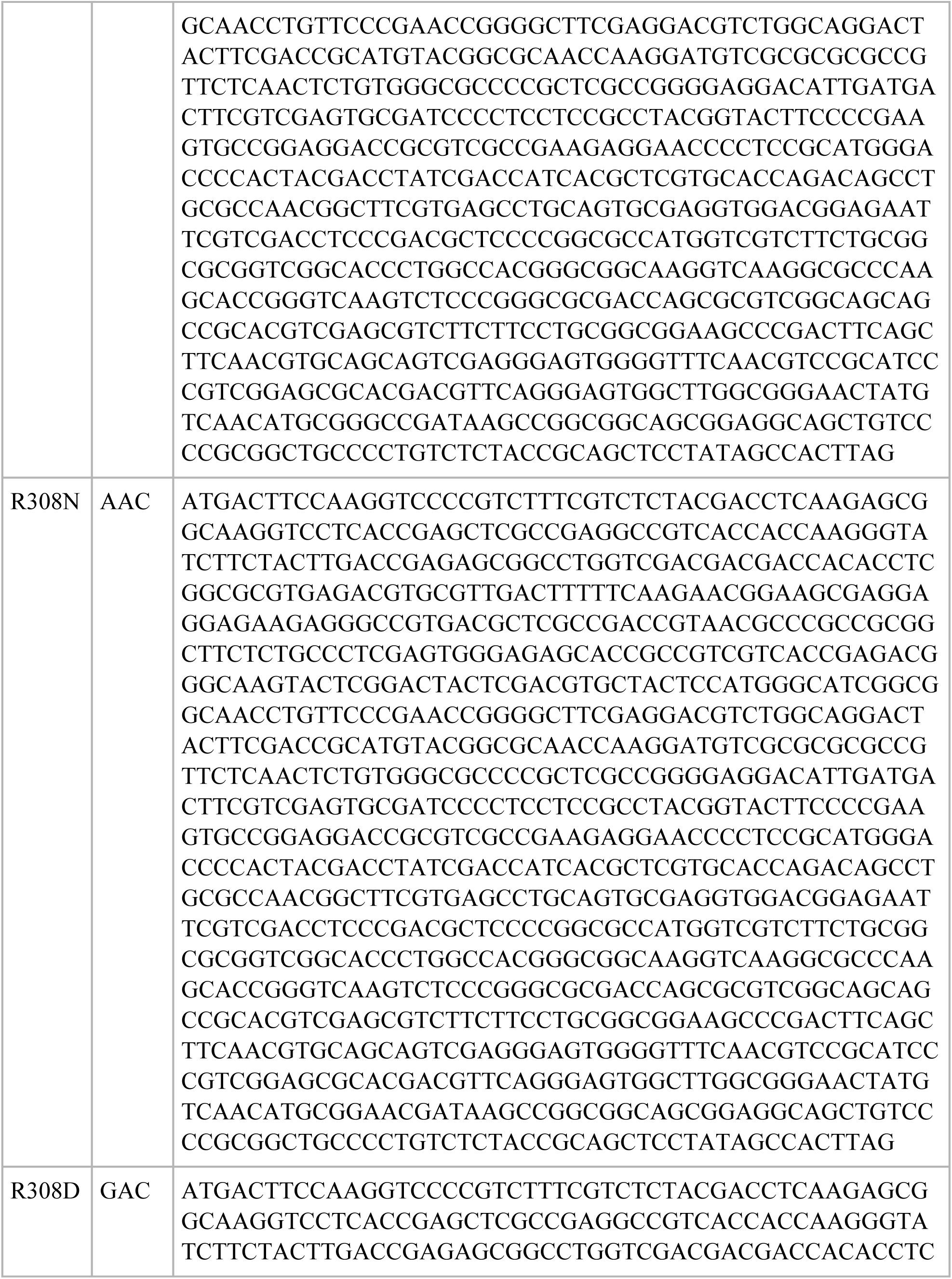
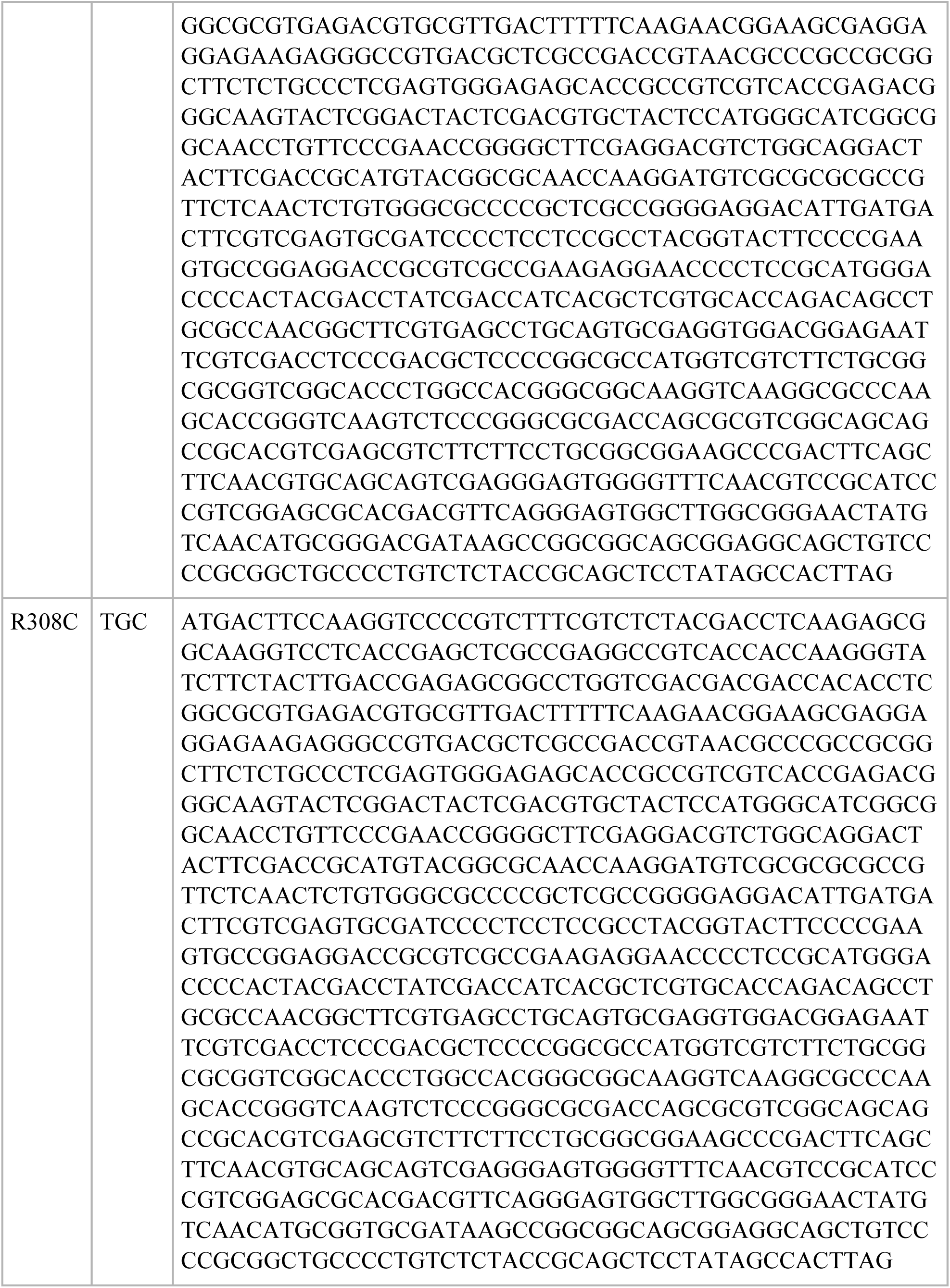
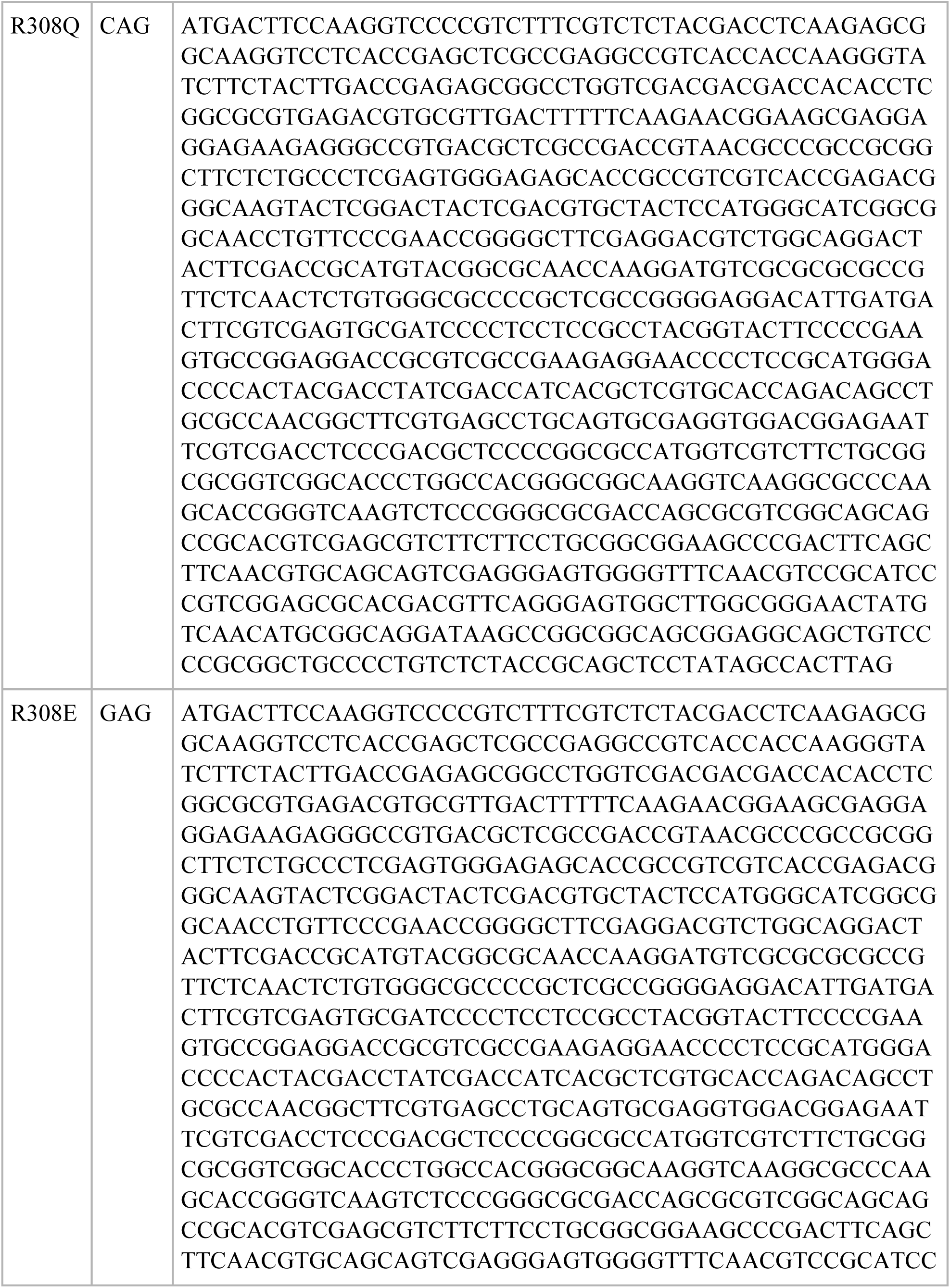
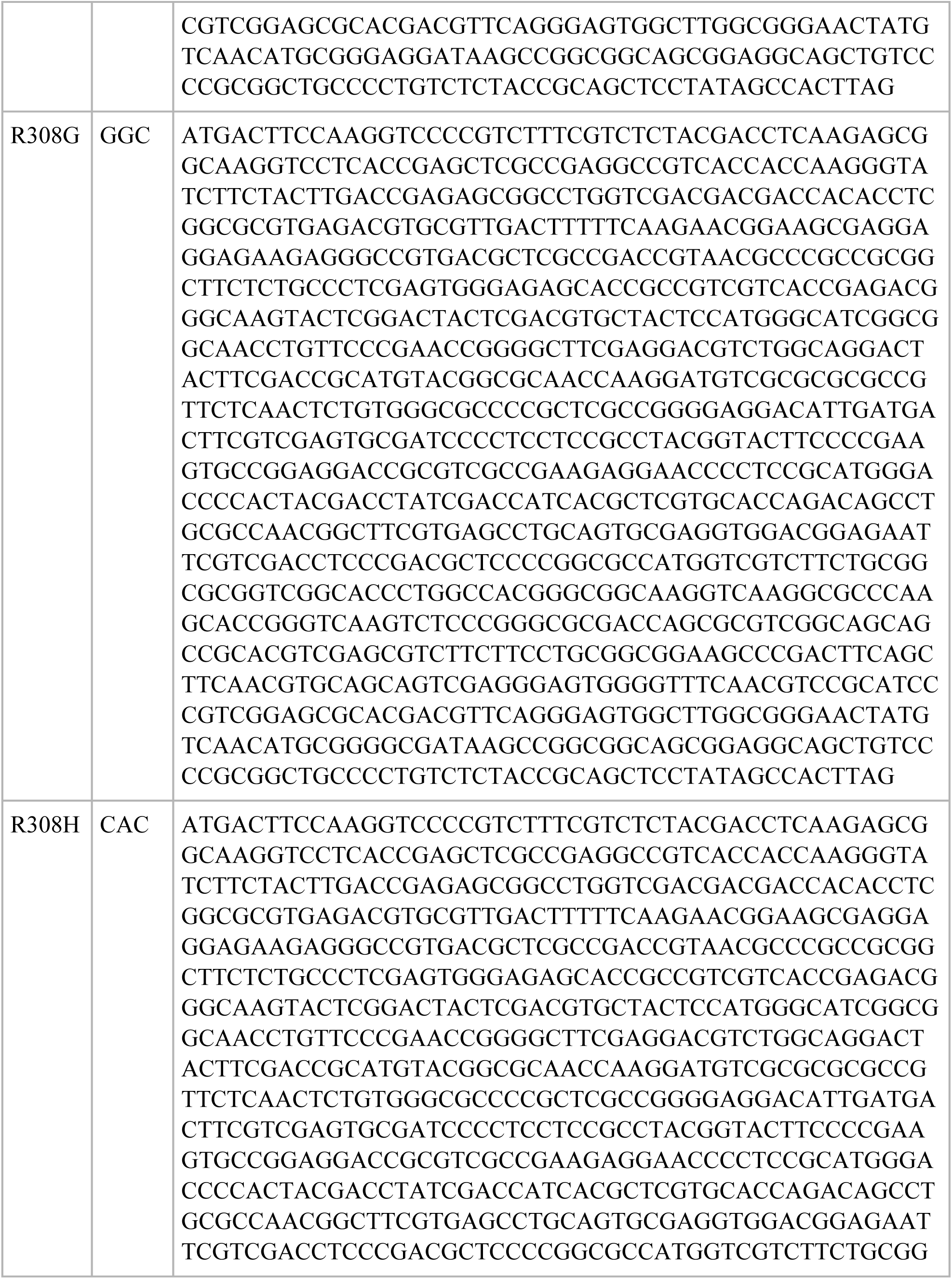
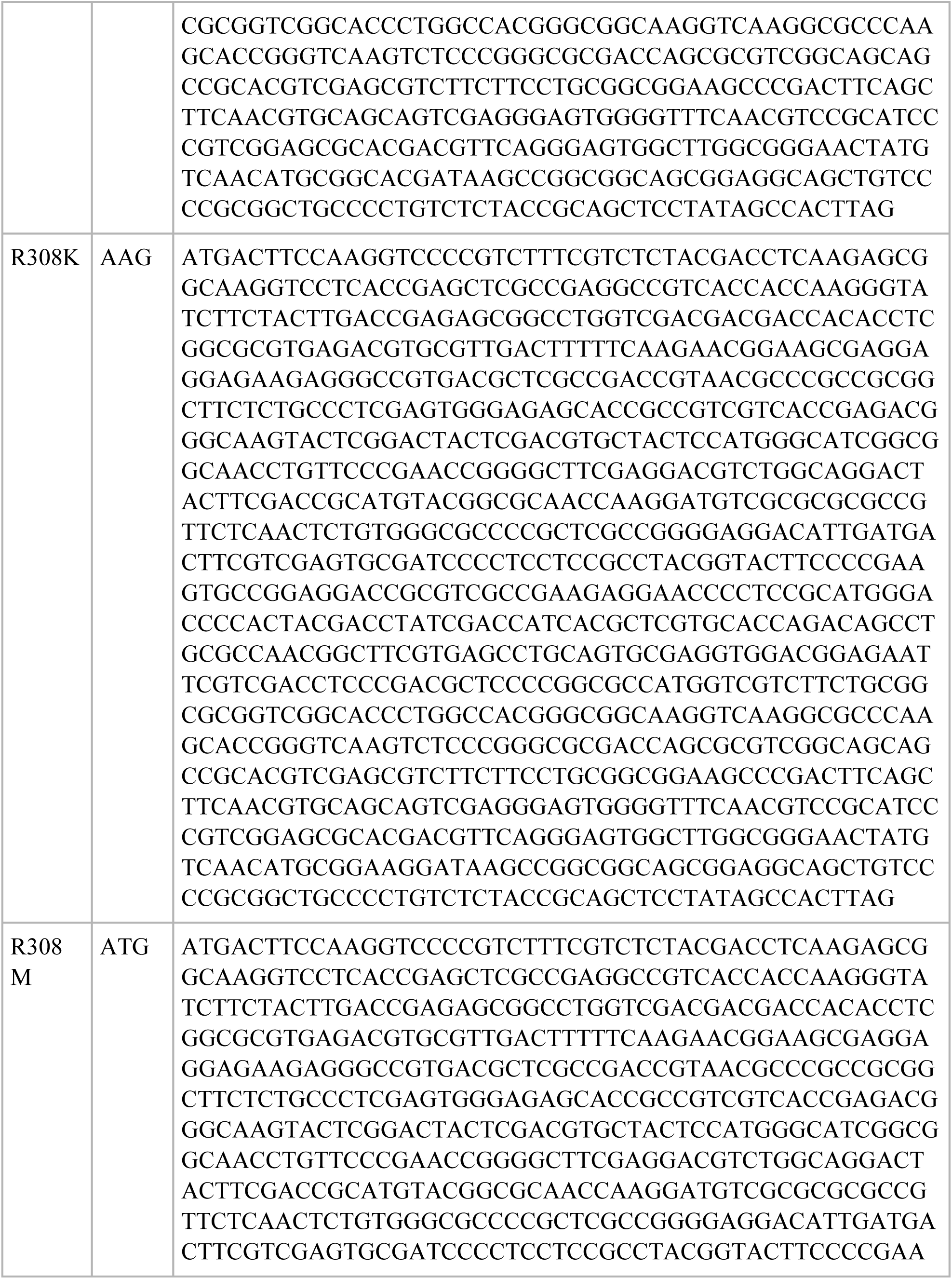
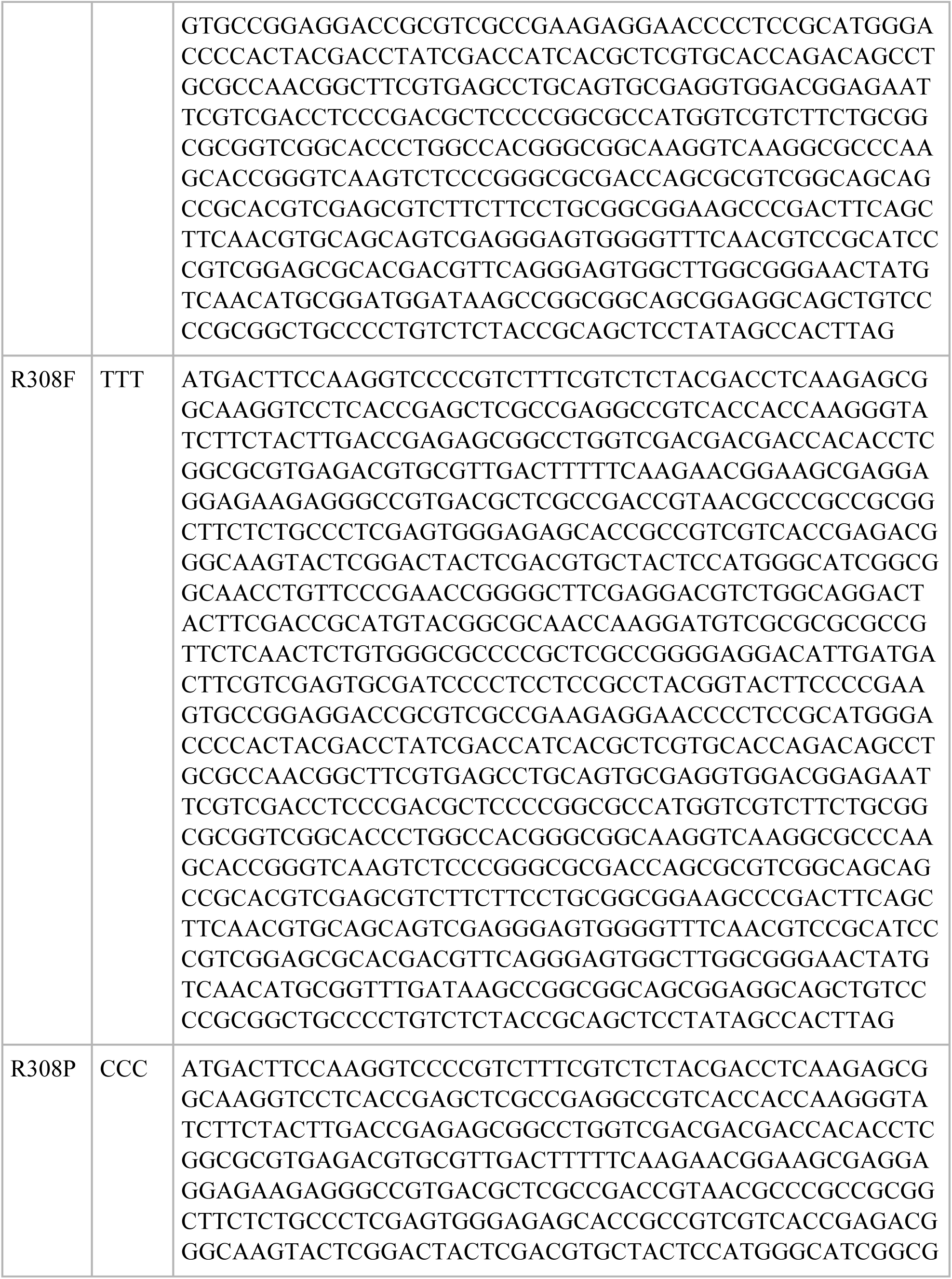
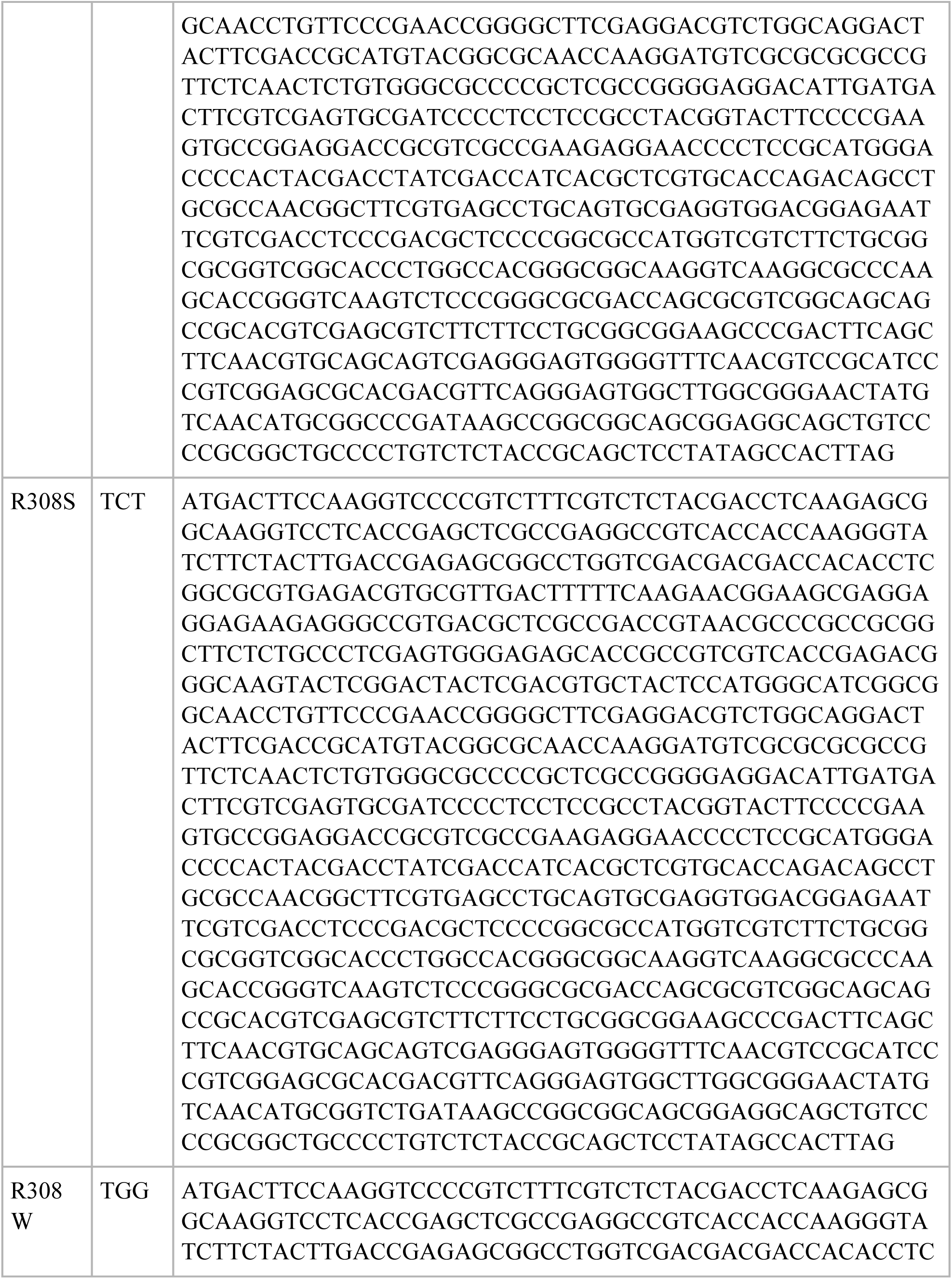
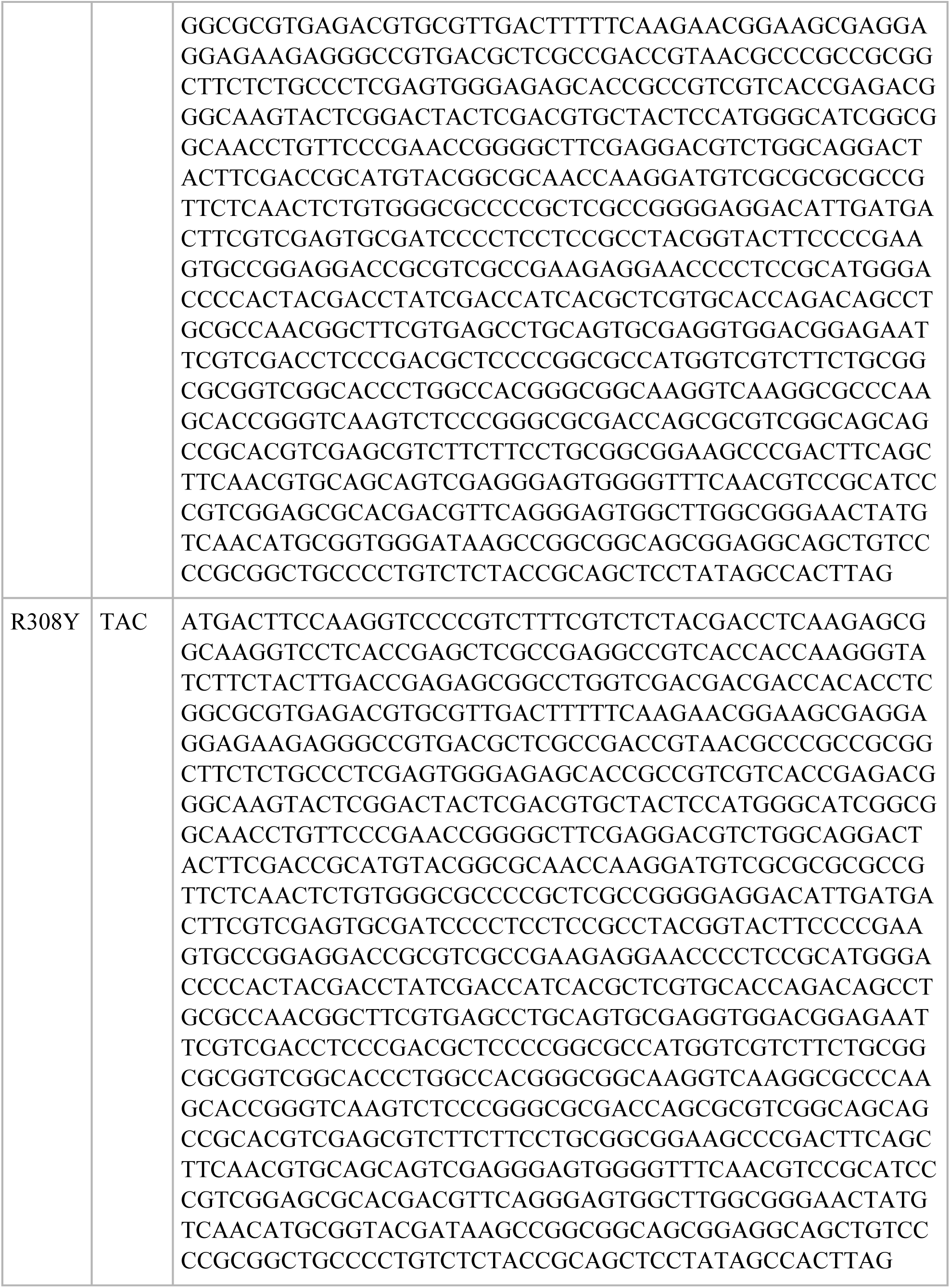
Mutant sequences of acDAOC/DACS.

